# Extracellular salicylic acid activates immune signaling through cell-surface receptors

**DOI:** 10.64898/2026.06.04.730219

**Authors:** Qi Li, Mingxi Zhou, Chetna Sharma, Peter A. Ramdhan, Cassidy R. Louwerse, Benjamin A. Merritt, Fahong Yu, Yanping Zhang, Chenglong Li, Xin Wang, Zhonglin Mou

## Abstract

Salicylic acid (SA) is a central immune hormone that accumulates in both intracellular and extracellular compartments during pathogen infection. While intracellular SA signaling is well established, whether extracellular SA (eSA) directly activates immune responses remains unknown. Here we show that eSA functions as an extracellular signal potentially perceived by the plasma membrane-localized lectin receptor kinases LecRK-I.8 and LecRK-VI.2 in *Arabidopsis*. The extracellular domains of both receptors bind SA with micromolar affinity, SA rapidly induces LecRK-I.8 phosphorylation, and the kinase activities of LecRK-I.8 and LecRK-VI.2 are required for downstream signaling. Moreover, mutagenesis of a computationally predicted binding pocket in LecRK-VI.2 abolishes SA binding and immune function, providing evidence for receptor-mediated perception. Genetic analyses further demonstrate that LecRK receptors are required for SA-induced resistance, transcriptional reprogramming, and phosphoproteomic responses. Together, these findings expand current models of SA signaling by revealing a potential receptor-mediated cell-surface perception mechanism for a classical immune hormone.

---

Over the past 50 years, salicylic acid (SA) has emerged as a central regulator of plant immunity and stress responses (*1*). It enhances tolerance to abiotic stresses such as drought, heat, UV radiation, and heavy metals, and regulates key developmental processes like germination, flowering, and senescence (*2*). Most critically, SA orchestrates multiple layers of plant immunity, including pattern-triggered immunity (PTI), effector-triggered immunity (ETI), and systemic acquired resistance (SAR) through its biosynthesis and signaling (*1, 3*). PTI is initiated by cell-surface recognition of conserved microbe-associated molecular patterns, restricting pathogen colonization (*4*). To overcome PTI, pathogens deploy effectors that suppress host defenses, leaving only basal resistance in susceptible hosts (*5*). In resistant hosts, pathogen-derived effectors are directly or indirectly recognized by intracellular nucleotide-binding leucine-rich repeat proteins, triggering ETI (*5*). Activation of PTI and ETI generates mobile immune signals that travel to distal tissues and promote the accumulation of extracellular NAD(P) [eNAD(P)] (*6, 7*). eNAD(P) is subsequently perceived by a group of plasma membrane-localized legume-like (L-type) lectin receptor kinases (LecRKs), which in turn activate downstream SA signaling to establish SAR (*8–10*).

In plants, SA signaling is mediated primarily by NONEXPRESSOR OF PATHOGENESIS-RELATED GENES (NPR) proteins, which function as SA receptors and transcriptional coactivators or corepressors that reprogram immune gene expression (*11–16*). Upon pathogen infection, SA accumulates in both intracellular and extracellular compartments (*17–19*). Whereas intracellular SA (iSA) signaling pathways are well understood (*1, 20*), the role of extracellular SA (eSA) remains poorly defined. In the apoplast, eSA has been proposed to function either as a SAR mobile signal or as a passive antimicrobial compound contributing to age-related resistance (ARR) (*17, 19, 21*). Genetic evidence further suggests that SA is actively transported by ATP-binding cassette transporters PLEIOTROPIC DRUG RESISTANCE8 (PDR8) and PDR12 to the apoplast and that iSA alone is insufficient to fully confer ARR (*22*). However, whether eSA directly triggers immune signaling at the cell surface remains unclear.

Apoplastic washing fluid (AWF) assays suggest that eSA levels reach 40-100 μM after pathogen infection (*23*). Because SA is not uniformly distributed, AWF measurements likely underestimate local eSA concentrations. Indeed, *in situ* measurements using an *Acinetobacter* sp. ADP1 SA biosensor indicate that eSA levels can reach ∼0.38 mM (*24*), placing physiologically relevant concentrations in the high micromolar to sub-millimolar range. Consistent with these estimates, such concentrations are commonly used to interrogate SA signaling mechanisms in plant immunity (*16, 25, 26*).

To determine whether eSA functions as an active immune signal, we depleted eSA in *Arabidopsis thaliana* (hereafter *Arabidopsis*) and systematically dissected its role in immune activation. We further identify LecRK-I.8 and LecRK-VI.2 as potential eSA receptors that directly bind SA and are required for SA-mediated immune signaling. These findings support a model in which plants sense eSA at the plasma membrane to trigger both local and systemic immune responses.

## SA induces LecRK-dependent resistance

Our previous work has shown that eNAD(P) is a critical SAR signal that functions downstream of a signal amplification loop in systemic tissues (*7*). While eNAD(P) induces SAR via SA signaling, SA itself also contributes to SAR signal amplification (*6*), raising the question of whether SA signaling depends on eNAD(P). To address this question, we tested SA-induced resistance to the bacterial pathogen *Pseudomonas syringae* pv. *maculicola* ES4326 (*Psm*) in *fin4-3* (*27*), a mutant with significantly reduced eNAD(P) accumulation (*7*), and *35S:CD38* transgenic plants, where eNAD(P) is partially depleted (*28*). We used 0.5 mM SA, close to the upper physiological range, to robustly activate downstream signaling. SA treatment conferred similar levels of resistance in Col-0 (wild type), *fin4-3*, and *35S:CD38* plants (fig. S1, A to D), indicating that SA induces eNAD(P)-independent resistance. We next examined previously characterized eNAD(P) receptor mutants: three T-DNA insertion alleles of *LecRK-VI.2* (*9*), *lecrk-I.8-2* (a T-DNA insertion line) (*8*), *lecrk-VI* (CRISPR/Cas9 knockout of clade VI members) (*10*), and *lecrk-I.8-2/VI* (*lecrk-I.8-2 lecrk-VI*) (*10*). Surprisingly, SA-induced resistance was significantly attenuated in these mutants (fig. S1, E to G), suggesting that these LecRKs function downstream of SA, in addition to serving as eNAD(P) receptors.

To confirm the phenotype of the T-DNA insertion mutant *lecrk-I.8-2*, we generated two independent deletion alleles of *LecRK-I.8*, *lecrk-I.8-cr* lines #2 and #3, using CRISPR/Cas9 with gRNAs targeting both the 5’ and 3’ ends of the gene (fig. S2A). Unexpectedly, these new alleles exhibited no defects in either NAD^+^-or SA-induced resistance (fig. S2, B to D), suggesting that *lecrk-I.8-2* may be a dominant-negative (DN) mutant. Indeed, overexpression of the truncated *lecrk-I.8-2DN* fragment markedly suppressed both NAD^+^-and SA-induced resistance in Col-0, *lecrk-VI*, and *lecrk-I.8-2/VI* backgrounds (fig. S2, E to M), supporting a role of LecRK-I.8 in eNAD(P)-and SA-mediated signaling and highlighting functional redundancy among LecRKs.

The *Arabidopsis* genome encodes 43 full-length L-type LecRKs, grouped into nine clades and five singletons (*29*). We used CRISPR/Cas9 to generate knockouts for the 43 *LecRKs*, grouped by clades or gene clusters (fig. S3, A to K), to systemically and unbiasedly investigate their roles in plant immunity. The CRISPR/Cas9-generated mutants exhibited normal morphology except for the *leck-S.1/4/6/7* knockout (targeting *LecRK-S.1*, *-S.4*, -*S.6*, and *-S.7*), which was slightly smaller than Col-0 plants (fig. S4A). Among the higher-order mutants created (Table S1), only *lecrk-I.7-11* (knockout of clade I members 7-11) and the previously generated *lecrk-VI* (*10*) exhibited reduced SA-and NAD^+^-induced resistance, impaired SAR induction, and compromised basal resistance (Fig. 1A and fig. S4, B to D). This confirms that clade I and clade VI LecRKs are required for both eNAD(P)-and SA-mediated signaling. Interestingly, the *leck-S.1/4/6/7* mutant showed heightened resistance to *Psm* (Fig. 1A and fig. S4, B to D), suggesting that these singleton LecRKs may function as negative regulators of plant immunity. Furthermore, the *lecrk-IX* mutant exhibited reduced SAR induction and basal resistance (fig. S4, C and D), supporting that clade IX LecRKs play positive roles in plant immunity (*30, 31*). By crossing *lecrk-I.1-6* or *lecrk-I.7-11* with *lecrk-VI*, we uncovered functional redundancy between clade I and VI LecRKs. SA-induced resistance was significantly reduced in *lecrk-I.7-11/VI*, but not in *lecrk-I.1-6/VI*, compared to *lecrk-VI* (Fig. 1B). Furthermore, *lecrk-I/VI*, a cross between *lecrk-I.1-6/VI* and *lecrk-I.7-11/VI*, which disrupted all 11 clade I and 4 clade VI members, resembled *lecrk-I.7-11/VI* (Fig. 1B), suggesting that LecRK-I.7-11 and clade VI LecRKs have partially redundant roles in SA-mediated immunity. Since LecRK-I.8 and clade VI LecRKs form receptor complexes with BRASSINOSTEROID INSENSITIVE1-ASSOCIATED RECEPTOR KINASE1 (BAK1) (*10*), we also tested SA-induced resistance in the *bak1-5 bkk1* double mutant (*32*), which is defective in BAK1-mediated immunity and its closest homolog BAK1-LIKE 1 (BKK1). Notably, SA-induced resistance to *Psm* was significantly impaired in *bak1-5 bkk1* (Fig. 1C), indicating that SA-triggered resistance depends on the LecRK-BAK1/BKK1 receptor complex.

**Fig. 1.**
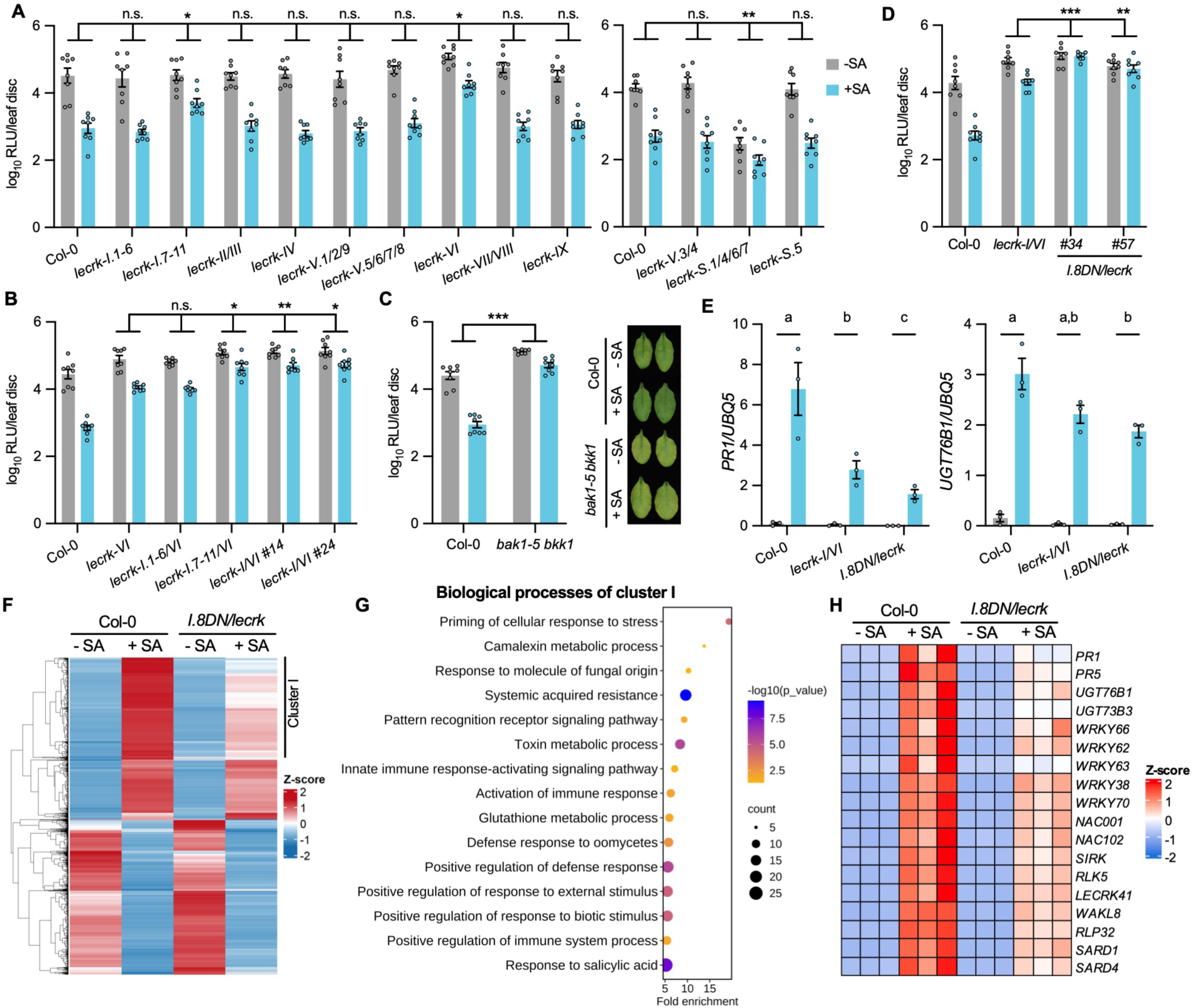
Clade I/VI LecRKs and BAK1 are essential for SA-induced immune signaling. (**A** to **D**) SA-induced resistance to *Psm* in CRISPR/Cas9-generated higher-order *lecrk* mutants (**A** and **B**), *bak1-5 bkk1*(**C**), and *I.8DN/lecrk* (**D**). Leaves on 4-week-old plants were infiltrated with water (-SA) or 0.5 mM SA (+SA). Four h later, the infiltrated leaves were inoculated with *Psm_lux* (OD_600_ = 0.001). Bacterial populations were measured and disease symptoms were photographed at 2.5 d post-inoculation (dpi). Bars represent means ± standard error of the mean (SEM) (n = 8). Asterisks denote significant differences (\*\*\**P* < 0.001, \*\**P* < 0.01, \**P* < 0.05, n.s., not significant; two-way ANOVA). RLU, relative light unit. (**E**) SA-induced expression of *PR1* and *UGT76B1* in *lecrk-I/VI* and *I.8DN/lecrk* plants. Leaves on 4-week-old plants were infiltrated with water or 0.5 mM SA and collected 4 h later for qPCR analysis. Bars represent means ± SEM (n = 3). Different letters denote significant differences (*P* < 0.05; two-way ANOVA). (**F**) Heatmap of differentially expressed genes (DEGs) in Col-0 and *I.8DN/lecrk* treated with water or 0.5 mM SA for 4 h. DEGs were defined as genes with an adjusted *P* value < 0.05 and |log₂(fold change)| > 2 relative to water treatment. Gene expression levels (RPKM) were obtained from RNA-seq data and normalized using z-scores. Rows represent individual genes and columns represent genotypes/treatments. The expression level is the average of three independent biological samples. Genes were grouped by hierarchical clustering based on expression patterns. The color scale reflects relative expression levels (z-scores). (**G**) Gene Ontology (GO) enrichment analysis of DEGs in cluster I shown in (**F**). The top 15 most significantly enriched biological process terms are displayed. GO analysis was performed using the PANTHER classification system. (**H**) Heatmap showing expression of selected defense-related genes in Col-0 and *I.8DN/lecrk* treated with water or 0.5 mM SA for 4 h. Gene expression levels (RPKM) were obtained from RNA-seq data and normalized using z-scores. The color scale represents relative expression levels (z-scores).

Consistent with their compromised SA-induced immunity, RNA sequencing (RNA-seq) analysis revealed that SA-triggered transcriptome reprogramming was attenuated in both *lecrk-I.7-11/VI* and *bak1-5 bkk1*, with the effect more pronounced in *lecrk-I.7-11/VI* (fig. S4, E to G and Data S1). We found that *pUBQ10:lecrk-I.8-2DN lecrk-I.8-2/VI* (hereafter *I.8DN/lecrk*) lines exhibited even more dramatic defects in SA-induced disease resistance and marker gene expression than *lecrk-I/VI* (Fig. 1, D and E). SA-induced transcriptomic changes were also markedly attenuated in *I.8DN/lecrk* (line #34) plants (Fig. 1F and Data S2). In particular, the activation of SA-related immune pathways and SA-responsive genes was significantly reduced (Fig. 1, G and H), indicating impaired SA signaling. Taken together, our findings demonstrate that LecRKs, mainly from clade I and clade VI, play essential and partially redundant roles in SA-mediated immune signaling.

### eSA contributes to basal immunity and SAR

To directly define the role of eSA, we generated transgenic *35S:eNahG-GFP* plants (hereafter *eNahG*), in which the PR1 signal peptide directs secretion of the salicylate hydroxylase NahG to the apoplast (fig. S5A). The eNahG-GFP protein accumulated in the apoplast without affecting plant morphology (fig. S5, B and C). Following *Psm* infection, SA accumulation in local tissues of *eNahG* plants was reduced to ∼60% of wild-type levels, whereas it was nearly abolished in *35S:NahG* plants (hereafter *NahG*), in which *NahG* is expressed intracellularly (Fig. 2A). In systemic tissues, SA levels did not increase in *NahG*, but were modestly elevated in *eNahG*, (Fig. 2B). Notably, apoplastic SA was nearly completely depleted in both local and systemic tissues of *eNahG* plants (Fig. 2, C and D), whereas iSA was largely preserved in local tissues and only partially reduced in systemic tissues (Fig. 2, E and F). These results indicate that eNahG preferentially depletes eSA while largely maintaining iSA, particularly in local tissues.

**Fig. 2.**
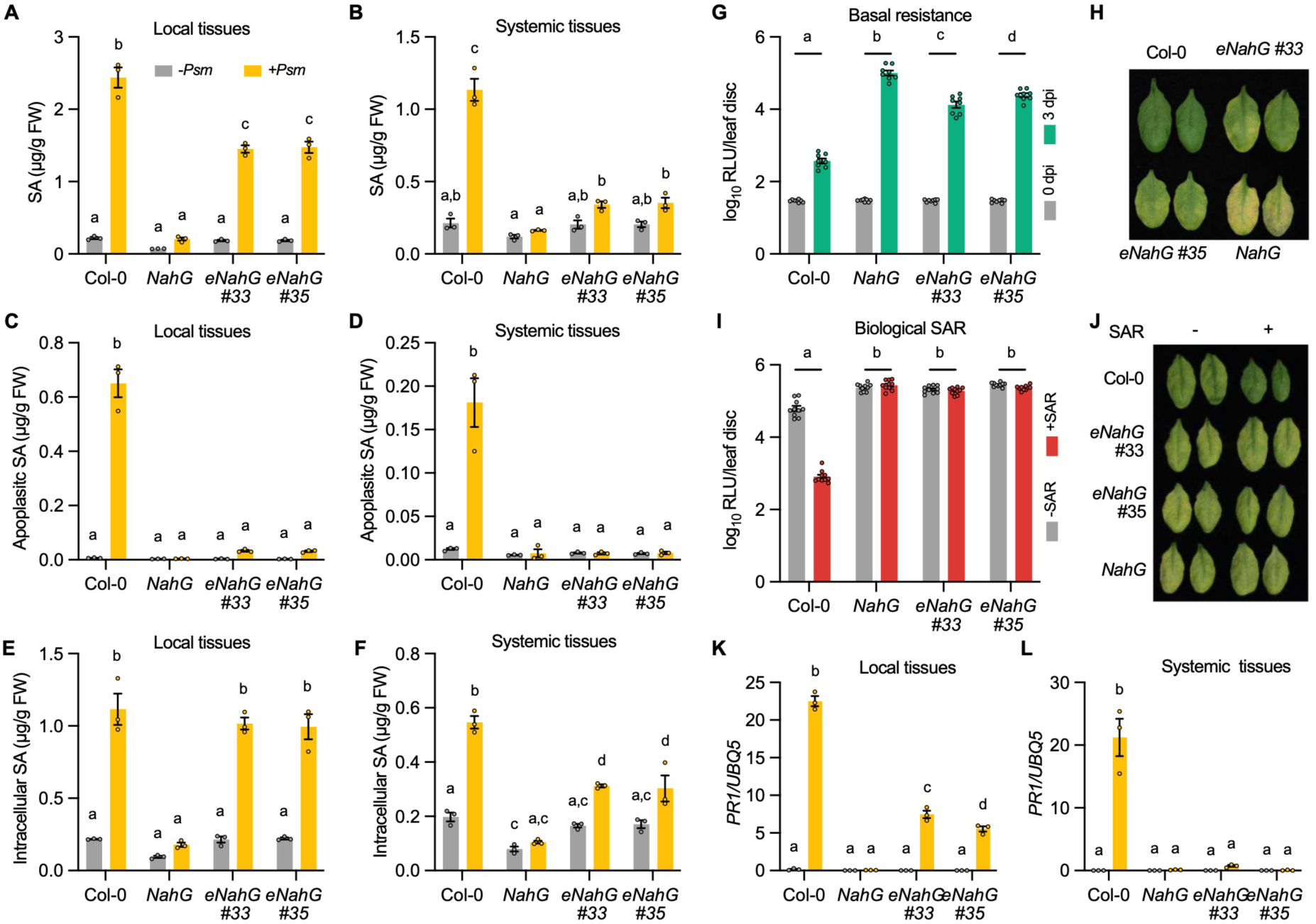
Extracellular SA is critical for activating local defenses and indispensable for systemic immune signaling. (**A** to **F**) *Psm*-induced accumulation of total (apoplastic + intracellular) SA (**A** and **B**), apoplastic SA (**C** and **D**), and intracellular SA (**E** and **F**) levels in local (**A**, **C**, and **E**) and systemic (**B**, **D**, and **F**) tissues of Col-0, *NahG*, and *eNahG* plants. For local tissues, leaves on 4-week-old plants were infiltrated with 1 mM MgCl₂ (-*Psm*) or *Psm* (OD_600_ = 0.001, +*Psm*), and the infiltrated leaves were collected 24 h later. For systemic tissues, leaves on 4-week-old plants were infiltrated with 1 mM MgCl₂ or *Psm* (OD_600_ = 0.004), and the upper, untreated leaves were collected 48 h later. Total, apoplastic, and intracellular SA were subsequently measured. Bars represent means ± SEM (n = 3). Different letters denote significant differences (*P* < 0.05; one-way ANOVA). (**G** and **H**) Basal resistance of Col-0, *NahG*, and *eNahG* plants. Leaves on 4-week-old plants were inoculated with *Psm_lux* (OD_600_ = 0.0002), and bacterial populations were measured at 0 and 3 dpi. Disease symptoms (**H**) were photographed at 3 dpi. (**I** and **J**) Biological induction of SAR in Col-0, *NahG*, and *eNahG* plants. Three lower leaves on each 4-week-old plant were infiltrated with 1 mM MgCl_2_ (-SAR) or *Psm* (OD_600_ = 0.004, +SAR). Two d later, one systemic leaf on each plant was inoculated with *Psm_lux* (OD_600_ = 0.001). Bacterial populations in the systemic leaves were measured at 2.5 dpi. Disease symptoms (**J**) were photographed at 2.5 dpi. In (**G** and **I**), bars represent means ± SEM (n = 8), and different letters denote significant differences (*P* < 0.05; two-way ANOVA). **(K** and **L)** *Psm*-induced *PR1* expression in local (**K**) and systemic (**L**) tissues of Col-0, *NahG*, and *eNahG* plants. Local and systemic tissues were prepared as in (**A** to **F**). Gene expression was analyzed by qPCR. Bars represent means ± SEM (n = 3). Different letters denote significant differences (*P* < 0.05; one-way ANOVA).

We next assessed the contribution of eSA to plant immunity. Basal resistance to *Psm* was significantly reduced in *eNahG* plants, although less severely than in *NahG* plants (Fig. 2, G and H). Moreover, SAR was completely abolished in both *NahG* and *eNahG* plants (Fig. 2, I and J), indicating that eSA is required for full basal immunity and SAR.

Consistent with these findings, infiltration of purified NahG protein into the apoplast following *P. syringae* pv. *tomato* DC3000 (*Pst*) infection increases susceptibility (*17*), reflecting the impact of selective eSA depletion on immunity. Similarly, co-infiltration of purified NahG protein with *Psm*, followed by re-infiltration at 12 h post-inoculation, resulted in >20-fold increased bacterial growth (fig. S5D). In addition, *svp* mutants, which exhibit reduced eSA while maintaining near wild-type iSA levels showed enhanced susceptibility and reduced defense gene expression (fig. S5, E to H) (*23*). Together, these observations support a critical and distinct role for eSA in plant immunity.

Although a small fraction of NahG may remain intracellular in *eNahG* plants, iSA levels were only minimally affected following infection, indicating that iSA signaling is largely preserved. Consistent with this, NAD^+^-induced local resistance was abolished in *NahG* but remained comparable between Col-0 and *eNahG* (fig. S6A), and NAD^+^-induced systemic resistance was impaired in *NahG* but partially retained in *eNahG* (fig. S6B). Given that NAD^+^ is membrane-impermeable (*33, 34*), these results indicate that iSA signaling remains functional in *eNahG* plants and operates downstream of LecRKs receptors.

To further define the signaling role of eSA, we performed RNA-seq analysis of Col-0, *NahG*, and *eNahG* plants following *Psm* infection. Compared to Col-0, pathogen-induced transcriptional reprogramming was strongly attenuated in *NahG* and partially reduced in *eNahG*, particularly for immune-related genes (fig. S7, A to F and Data S3). For example, genes in cluster 5, enriched for immune responses functions, were strongly induced in Col-0 but not in *NahG*, with intermediate induction in *eNahG* (fig. S7, B, C and F). In contrast, cluster 1 genes, associated with hypoxia responses, were similarly reduced in both *NahG* and *eNahG* (fig. S7, B, C and D), suggesting a potential role for eSA in regulating hypoxia-related pathways. These results were validated by qPCR analysis of defense-related genes, which showed intermediate expression in *eNahG* relative to Col-0 and *NahG* in local tissues (Fig. 2K and fig. S7G), but *NahG*-like expression in systemic tissues (Fig. 2L and fig. S7H). Collectively, these findings support the idea that eSA contributes to full activation of both local immune responses and SAR.

### LecRK-I.8 and LecRK-VI.2 directly bind SA

To determine whether eSA functions as an active signaling molecule in the apoplast, we investigated whether LecRK-I.8 and LecRK-VI.2 directly bind SA. To this end, the extracellular domains of LecRK-I.8 and LecRK-VI.2 (eLecRK-I.8 and eLecRK-VI.2) were expressed in *Nicotiana benthamiana* and purified from the AWF (fig. S8A). We first conducted binding assays using [^3^H]-SA. The total SA-binding activities of eLecRK-I.8 and eLecRK-VI.2 were significantly higher than those of the no protein control (Fig. 3A). Furthermore, [^3^H]-SA binding to eLecRK-I.8 and eLecRK-VI.2 was effectively outcompeted by unlabeled SA (Fig. 3A), consistent with specific binding.

**Fig. 3.**
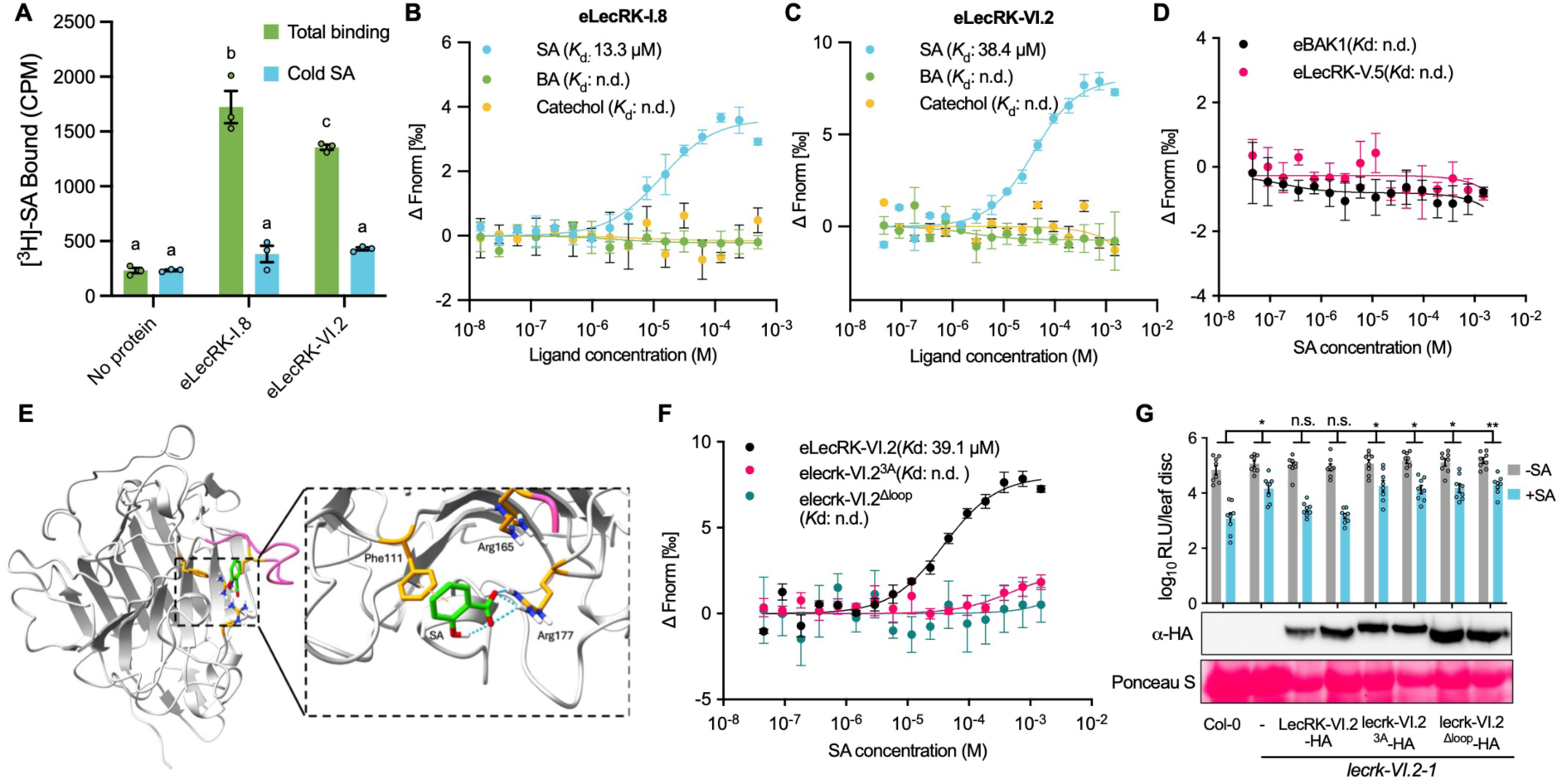
SA binds to the extracellular domains of LecRK-I.8 and LecRK-VI.2. (**A**) Binding of [³H]-SA to recombinant eLecRK-I.8 and eLecRK-VI.2 proteins. No-protein samples served as the negative control. CPM, count per minute. Bars represent means ± SEM (n = 3). Different letters denote significant differences (*P* < 0.05; one-way ANOVA). (**B** and **C**) MST analysis of binding between eLecRK-I.8 (**B**) or eLecRK-VI.2 (**C**) and SA or its analogs. MST was performed using fluorescently labeled proteins and titrated concentrations of SA, catechol, and BA. Protein and ligand concentrations are described in Methods. ΔFnorm represents the normalized fluorescence changes of labeled proteins. Data represent means ± SEM (n = 3 independent experiments). Dissociation constants (*K*_d_) were calculated by non-linear regression, and best-fit values are shown. n.d., not detected. (**D**) MST analysis of SA binding to eBAK1 or eLecRK-V.5. (**E**) Computational model of SA bound to the extracellular domain of eLecRK-VI.2. Hydrogen bonds (blue dashed lines), aromatic interactions (orange dashed line), and the loop targeted for deletion (pink) are indicated. (**F**) MST analysis of SA binding to wild-type eLecRK-VI.2 and the 3A and Δloop mutant variants. (**G**) SA-induced resistance depends on SA binding to the extracellular domain of LecRK-VI.2. Leaves on T_2_ transgenic *lecrk-VI.2-1* plants expressing LecRK-VI.2-HA, lecrk-VI.2^3A^-HA, or lecrk-VI.2^Δloop^-HA were infiltrated with water (-SA) or 0.5 mM SA (+SA) and inoculated 4 h later with *Psm_lux* (OD_600_ = 0.001). Bars represent means ± SEM (n = 8). Asterisks denote significant differences (\*\**P* < 0.01, \**P* < 0.05, n.s., not significant; two-way ANOVA). Protein levels were assessed by immunoblotting with Ponceau S staining of Rubisco as the loading control.

Next, microscale thermophoresis (MST) was used to confirm SA binding and determine the dissociation constant (*K*d). eLecRK-I.8 and eLecRK-VI.2 bound SA with *K*d values of 13.3 μM and 38.4 μM, respectively, in MST buffer (pH 5.7), whereas eLecRK-V.5 and eBAK1 showed no detectable binding (Fig. 3, B to D). Consistent with these affinities, SA concentrations as low as 50 μM significantly induced resistance to *Psm* and activated defense gene expression (fig. S8B) (*35*). Notably, measured eSA concentrations exceed these *K*d values (*23, 24*), indicating that physiological eSA levels are sufficient to achieve substantial receptor occupancy and activate downstream signaling.

Interestingly, eLecRK-I.8 also bound phosphatidylcholine (PC) with a *K*d value of 28.8 μM (fig. S8C), consistent with its role in perceiving PC species derived from insect eggs as egg-associated molecular patterns (*36*). In contrast, neither eLecRK-I.8 nor eLecRK-VI.2 bound the biologically inactive SA analogs catechol or benzoic acid (BA) in MST buffer (Fig. 3, B and C), indicating selective recognition of biologically relevant ligands rather than nonspecific binding. Furthermore, addition of eBAK1 did not alter the SA-binding affinities of eLecRK-I.8 and eLecRK-VI.2 (fig. S8, D and E), and SA did not enhance the association between LecRK-I.8 or LecRK-VI.2 with BAK1 (fig. S8F). Together, these results support a model in which LecRK-I.8 and LecRK-VI.2 function as eSA receptors that associate with BAK1 in preformed receptor complexes. Consistent with this role, the expression of *LecRK-I.8*, *LecRK-VI.2*, and several additional *LecRKs* was rapidly upregulated following SA treatment (fig. S8G).

Computational modeling of LecRK-SA interactions revealed a putative SA-binding pocket in the extracellular domain of LecRK-VI.2 but not in LecRK-I.8. We therefore focused on LecRK-VI.2. Fig. 3E illustrates several key contacts stabilizing SA binding. The SA phenolic hydroxyl forms an intramolecular hydrogen bond with its carboxylate group, while the carboxylate engages in multiple hydrogen bonds with Arg177, anchoring the ligand within the pocket. In addition, Phe111 participates in a T-shaped π-π interaction with the SA aromatic ring, further stabilizing the ligand binding. Consistent with this model, both a triple substitution mutant (F111A/R165A/R177A; 3A) and a loop-deletion mutant (lacking Val155-Ile166; Δloop) retained the predicted overall fold but lost SA binding activity and failed to complement the *lecrk-VI.2* defect in SA-induced *Psm* resistance (Fig. 3, F and G and fig. S8, H and I). These findings demonstrate that direct SA binding to LecRK-VI.2 is required for receptor function and downstream signaling.

### SA triggers LecRK-dependent phosphorylation

Both LecRK-I.8 and LecRK-VI.2 are active kinases (*8, 37*), prompting us to investigate whether SA signaling depends on their kinase activity. Treatment with the broad-spectrum kinase inhibitor K252a reduced SA-induced expression of defense genes and compromised SA-induced resistance to *Psm* (fig. S9, A to C), suggesting that kinase activity is essential for SA signaling. To determine whether SA activates the LecRK receptors via phosphorylation, we analyzed LecRK-I.8 and LecRK-VI.2 in transgenic *Arabidopsis* using Phos-tag gels. SA, but not BA, induced a mobility shift of LecRK-I.8 (Fig. 4A). The SA-triggered shift was detectable as early as 5 min, peaking at 30 min, and returning to basal levels by 90 min (Fig. 4B and fig. S9D), indicating rapid and transient receptor activation. We found that the SA-induced mobility shift of lecrk-I.8^K357E^-HA, a kinase-dead variant, was markedly reduced compared to wild-type LecRK-I.8-HA (Fig. 4C), suggesting that LecRK-I.8 activity contributed either directly or indirectly to the mobility shift observed in Phos-tag gels. We did not detect a mobility shift of LecRK-VI.2 under the same conditions (fig. S9E), possibly due to low phosphorylation levels or technical limitations. To evaluate the functional relevance of the receptor kinase activity *in vivo*, we introduced wild-type and kinase-dead forms of LecRK-I.8 and LecRK-VI.2 into their respective mutants. While LecRK-I.8-HA and LecRK-VI.2-HA restored SA-induced resistance, the kinase-dead variants (lecrk-I.8^K357E^-HA and lecrk-VI.2^D494N^-HA) failed to complement, despite comparable protein levels (Fig. 4, D and E and fig. S9, F and G), demonstrating the requirement of receptor kinase activity for SA-induced immunity.

**Fig. 4.**
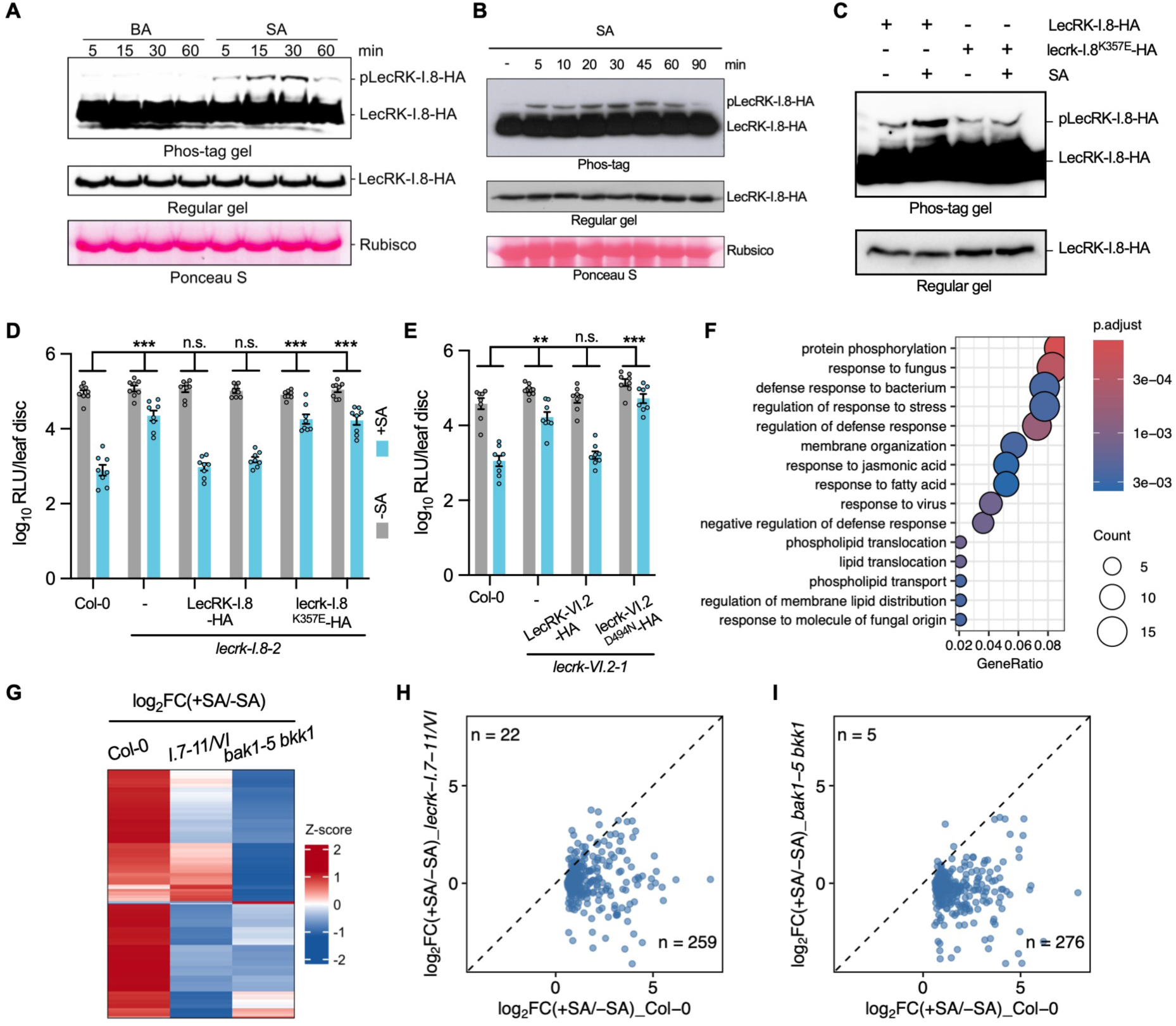
SA rapidly induces LecRK-I.8 phosphorylation and widespread LecRK-dependent phosphorylation changes. (**A**) Rapid induction of LecRK-I.8 phosphorylation by SA treatment. Leaves on 4-week-old *pUBQ10:LecRK-I.8-HA/lecrk-I.8-2* plants were infiltrated with 0.5 mM SA or BA (control). Total proteins were extracted from leaf tissues collected at the indicated time points and separated by Mn²⁺-Phos-tag or regular SDS-PAGE gels, followed by immunoblotting with anti-HA antibody. The upper shifted band (pLecRK-I.8-HA) indicates the phosphorylated form of LecRK-I.8-HA. Ponceau S staining of Rubisco was used as a loading control. (**B**) Rapid induction of LecRK-I.8 phosphorylation by SA treatment. Leaves on 4-week-old *pUBQ10:LecRK-I.8-HA/lecrk-I.8-2* plants were vacuum-infiltrated with 0.6 mM SA or water (-). Total proteins were extracted from leaf tissues collected at the indicated time points (the water-treated sample was collected at 20 min) and separated by Mn²⁺-Phos-tag or regular SDS-PAGE, followed by immunoblotting with anti-HA antibody. The upper shifted band (pLecRK-I.8-HA) indicates the phosphorylated form of LecRK-I.8-HA. (**C**) SA-induced phosphorylation of LecRK-I.8 depends on its kinase activity. Leaves on *pUBQ10:LecRK-I.8-HA/lecrk-I.8-2* and *pUBQ10:lecrk-I.8^K357E^-HA/lecrk-I.8-2* plants were infiltrated with water (-SA) or 0.5 mM SA. Total proteins were extracted 20 min later and analyzed as in (**A**). (**D** and **E**) SA-induced resistance depends on the kinase activity of LecRK-I.8 and LecRK-VI.2. Leaves on 4-week-old plants of the indicated genotypes were infiltrated with water (-SA) or 0.5 mM SA (+SA). Four h later, the same leaves were inoculated with *Psm_lux* (OD_600_ = 0.001). The bacterial populations in the leaves were measured at 2.5 dpi. Bars represent means ± SEM (n = 8). Asterisks denote significant differences (\*\*\**P* < 0.001, \*\**P* < 0.01, n.s., not significant; two-way ANOVA). (**F**) GO analysis of proteins that are rapidly phosphorylated by SA treatment in Col-0. The top 15 significantly enriched biological process terms are displayed. (**G** to **I**) Heatmap (**G**) and scatter plots (**H** and **I**) showing that most of the SA-upregulated phosphosites in Col-0 are less induced in *lecrk-I.7-11/VI* and *bak1-5 bkk1*. SA-upregulated phosphosites in Col-0 (*P* < 0.05, log_2_FC > 0.5) were retrieved from Col-0, *lecrk-I.7-11/VI*, and *bak1-5 bkk1* datasets and used for visualization. FC, fold change.

Given that SA induces both receptor phosphorylation and immune activation, we next performed phosphoproteomic profiling to assess SA-induced global protein phosphorylation in wild-type Col-0, *lecrk-I.7-11/VI*, and *bak1-5 bkk1*. Using a site localization probability cutoff of ≥ 0.75, a total of 8,246 unique phosphosites corresponding to 3,735 proteins were identified (Data S4). In Col-0, SA treatment rapidly triggered phosphorylation of proteins involved in protein phosphorylation, defense responses, and membrane organization (Fig. 4F and fig. S9H). Importantly, SA-induced global protein phosphorylation was largely abolished in *lecrk-I.7-11/VI* and *bak1-5 bkk1* plants (Fig. 4, G to I), underscoring the essential role of LecRKs and BAK1 in mediating SA-induced phosphorylation events. Together, these findings indicate that LecRKs mediate SA signaling through their kinase activity, likely by phosphorylating downstream targets involved in diverse cellular processes.

### SA induces LecRK-dependent early immune outputs

To determine whether SA directly activates receptor-mediated early signaling independently of transcription, we examined hallmark early immune responses (*38, 39*). SA and NAD⁺ did not elicit a detectable reactive oxygen species (ROS) burst in mature leaves or activate MAP kinases (MAPKs) in seedlings, whereas flg22 (a conserved 22-amino acid epitope from bacterial flagellin) elicited comparable ROS production and MAPK activation in Col-0, *NahG*, *eNahG*, and *lecrk-I.7-11/VI* plants (fig. S10, A to D). In contrast, SA and NAD⁺ induced weak but reproducible MAPK activation in mature leaves that was NPR1-independent and partially LecRK-dependent (fig. S10, E and F). Although neither treatment triggered Ca²⁺ influx in seedlings (fig. S10G), both elicited clear Ca²⁺ influx in mature leaves (Fig. 5, A and B); in protoplasts, 0.25 mM SA-induced Ca²⁺ influx was NPR1-independent and largely LecRK-dependent (Fig. 5C). Together, these results establish that LecRK receptors, but not NPR1, drive SA-triggered early signaling. Consistent with this, SA-induced stomatal closure was abolished in *lecrk-I/VI* and *I.8DN/lecrk* plants, whereas abscisic acid (ABA)-induced closure remained intact (Fig. 5, D and E), further supporting that LecRKs mediate SA-triggered early immune responses.

**Fig. 5.**
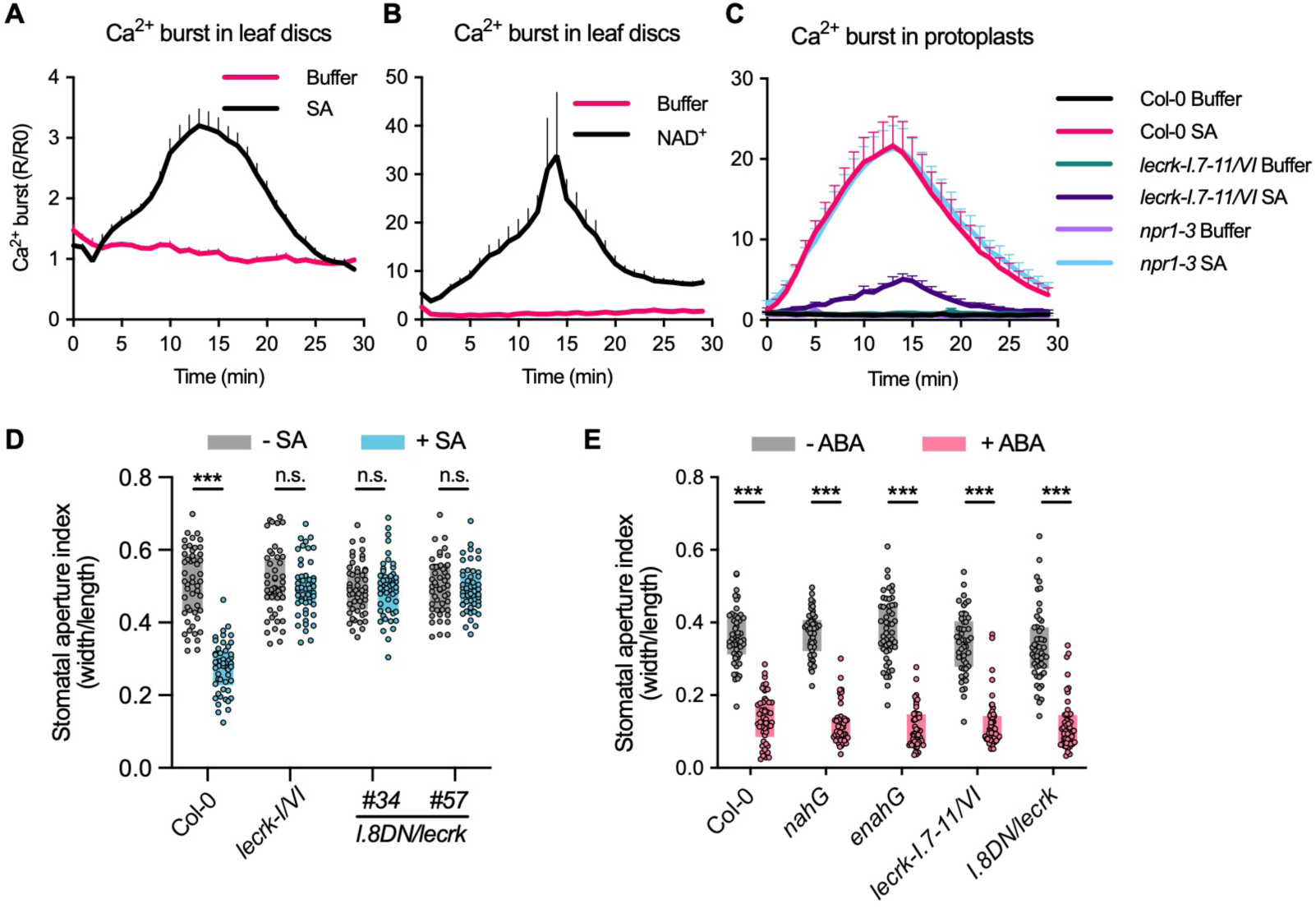
SA induces LecRK-dependent early immune responses. (**A** and **B**) SA-(**A**) and NAD^+^-triggered (**B**) cytosolic Ca^2+^ burst in adult plants. Leaf discs from 4-week-old cytAEQ plants were treated with buffer, or 0.5 mM SA or 0.5 mM NAD^+^ in the buffer. (**C**) SA-induced cytosolic Ca^2+^ burst in *lecrk-I.7-11/VI* and *npr1-3*. Protoplasts transiently expressing cytAEQ were treated with buffer or 0.25 mM SA in the buffer. In (**A** to **C**), Cytosolic Ca^2+^ burst was recorded as luminescence (R) immediately after treatment for indicated times, normalized to baseline luminescence before treatment (R/R₀). Data represent means ± SEM (n = 8). (**D** and **E)** SA-(**D**) and ABA-triggered (**E**) stomatal closure in the indicated genotypes. *Arabidopsis* epidermal peels were treated with mock (water for SA and 0.1% DMSO for ABA), SA (0.5 mM) (**D**), or ABA (5 μM) (**E**). Stomatal apertures were measured at 1 h post-treatment. Data are presented as box plots as described in Methods (n = 50). Statistical analysis was performed within each genotype (\*\*\**P* < 0.001; n.s., not significant; two-sided Student’s *t*-test).

## Discussion

Since SA was first shown to be both necessary and sufficient for SAR induction over three decades ago (*40, 41*), identifying its receptor has been a central goal in plant immunity research. Multiple forward genetic screens conducted in the early 1990s led to the discovery of NPR1, a master regulator of SA signaling, which was independently identified by several groups in 1997 (*11, 42, 43*). Subsequent studies revealed that NPR1 and its paralogs NPR3 and NPR4 directly bind SA (*12, 13*), and structural analyses confirmed the presence of *bona fide* SA binding domains (*15, 16*), establishing these proteins as iSA receptors. In parallel with genetic approaches, biochemical screens using radio-labeled SA or the photo-reactive SA analog 4-AzidoSA identified over 100 SA-binding proteins (SABPs) from protein extracts and microarrays (*44, 45*). These include key redox regulators such as catalase and ascorbate peroxidase (*44, 46*), as well as enzymes like carbonic anhydrase and methyl salicylate esterase, which play roles in stress responses and SAR signaling (*47, 48*). However, it remains unclear whether any of these SABPs function as genuine SA receptors, highlighting the ongoing challenge of distinguishing SABPs from true signal-transducing receptors.

Given their functional redundancy and plasma membrane localization, it is not surprising that LecRKs were not identified in earlier forward genetic or biochemical screens. Single *lecrk* mutants typically exhibit no or only mild SA-related phenotypes (*8*), making them difficult to detect in traditional genetic screens. Similarly, the low abundance and hydrophobic nature of transmembrane LecRKs hindered their recovery in biochemical assays. Our attention was drawn to LecRKs while investigating the interplay between SA and eNAD(P). We noticed that *lecrk-I.8-2* and *lecrk-VI.2*, previously characterized as eNAD(P) receptor mutants (*8, 9*), displayed slight but significant defects in SA-induced immunity. This observation motivated the development of *eNahG* plants, higher-order *lecrk* mutants, and dominant-negative *I.8DN/lecrk* lines to dissect the role of eSA in immunity and to evaluate the contribution of LecRKs to SA signaling. Moreover, computational modeling, combined with mutagenesis of the predicted SA-binding pocket, revealed that SA binding is critical for LecRK-VI.2 activation. Together, these complementary approaches provide compelling evidence that eSA functions as an essential extracellular immune signal and that LecRK-I.8 and LecRK-VI.2 act as eSA receptors, uncovering a potential cell-surface perception mechanism for SA.

While the interplay between cell-surface and intracellular SA receptors remains to be fully defined, our data reveal a tight functional coupling between extracellular and intracellular SA signaling. First, higher-order *lecrk* mutants and *I.8DN/lecrk* lines fail to mount SA-induced resistance even at upper physiological SA levels, suggesting that SA perception at the cell-surface is necessary. Consistently, *pdr8-4 pdr12-3*, a double mutant of the putative SA transporters PDR8 and PDR12, accumulates markedly reduced eSA but elevated iSA upon pathogen infection, yet exhibits compromised ARR and increased susceptibility to *Pst* relative to wild-type Col-0 (*22*). Similarly, *eNahG* plants remain susceptible despite near wild-type iSA levels. Together, these observations indicate that iSA alone is insufficient and support the idea that eSA perception is required for full immune activation. Second, eNAD^+^, an endogenous ligand of LecRK-I.8 and LecRK-VI.2, restores resistance in *eNahG* but not in *NahG* plants, placing LecRK function upstream of iSA. The stronger susceptibility of *NahG* compared to *eNahG* further suggests that iSA also contributes to an eSA-independent layer of immunity. Consistent with this model, LecRKs are required for SA-induced, NPR1-dependent transcriptional reprogramming, although the mechanistic link remains to be elucidated. Notably, SA also triggers rapid, NPR1-independent responses, including LecRK-dependent MAPK activation and Ca^2+^ influx, widespread protein phosphorylation, and inhibition of clathrin-mediated endocytosis and pollen tip growth (*49, 50*). Together, these findings suggest a dual-branch SA signaling framework in which LecRK-mediated extracellular perception both licenses iSA signaling and drives a rapid, non-transcriptional immune pathway.

While the evidence presented here strongly implicates LecRK-I.8 and LecRK-VI.2 in eSA perception, definitive classification of these proteins as *bona fide* eSA receptors will require further investigation. High-resolution structural studies, together with extensive structure-guided mutagenesis and biochemical analyses, are needed to establish the precise mode of SA recognition and determine how ligand binding activates receptor signaling. Such studies will provide a rigorous framework for understanding cell-surface SA perception and its integration with the well-established intracellular SA signaling pathways (*1, 20*).

## Materials and Methods

### Plant materials and growth conditions

*Arabidopsis thaliana* accession Columbia-0 (Col-0) was used as the wild type. All mutants and transgenic plants were in the Col-0 background. The following lines were previously described: *bak1-5 bkk1* (*32*), *lecrk-I.8-2* (*8*), *lecrk-VI.2-1* (*9*), *lecrk-VI.2-2* (*9*), *lecrk-VI.2-3* (*9*), *fin4-3* (*27*), *lecrk-VI* (*10*), *lecrk-I.8-2/VI* (*10*), *svp-31* (*52*), *svp-32* (*52*), *pLecRK-VI.2:LecRK-VI.2-HA/lecrk-VI.2-1* (*9*), *pLecRK-VI.2:lecrk-VI.2^D494N^-HA/lecrk-VI.2-1* (*9*), *35S:CD38* (*28*), *35S:NahG* (*53*) and Col-0 expressing cytosolic aequorin (cytAEQ) (*54*). *lecrk-I.1-6*, *lecrk-I.7-11*, *lecrk-II/III*, *lecrk-IV*, *lecrk-V.1/2/9*, *lecrk-V.3/4*, *lecrk-V.5/6/7/8*, *lecrk-VII/VIII*, *lecrk-IX*, *lecrk-S.1/4/6/7*, *lecrk-S.5*, and *lecrk-I.8-cr* were generated using CRSIPR/Cas9 in this study. *lecrk-I.1-6/VI*, *lecrk-I.7-11/VI*, and *lecrk-I/VI* were generated by cross in this study. *35S:eNahG-GFP*, *pUBQ10:LecRK-I.8-DN*/Col-0, *pUBQ10:LecRK-I.8-DN/lecrk-VI*, *pUBQ10:LecRK-I.8-DN/lecrk-I.8-2/VI*, *pUBQ10:LecRK-I.8-HA/lecrk-I.8-2*, *pUBQ10:lecrk-I.8^K357E^-HA/lecrk-I.8-2*, *pUBQ10:LecRK-VI.2/lecrk-VI.2-1*, *pUBQ10:lecrk-VI.2^3A^/lecrk-VI.2-1*, and *pUBQ10:lecrk-VI.2^Δloop^/lecrk-VI.2-1* were generated by transformation in this study. *Arabidopsis* plants were grown in soil (Berger BM2 Seed Germination Mix supplemented with fertilizer) in a controlled growth room at 24/22 °C (day/night) under a 12/12 h light/dark photoperiod, 50-60% relative humidity, and 100 μE m^-2^ s^-1^ light intensity. Four-week-old *Arabidopsis* plants were used for all experiments unless otherwise specified. *Nicotiana benthamiana* plants were grown in soil in a growth room with the same conditions. Five-week-old *N. benthamiana* was used for recombinant protein expression. For *Arabidopsis* seedling growth on Murashige and Skoog (MS) medium, seeds were surface sterilized with 15% (v/v) bleach, washed three times with sterile water, and germinated on sterile half-strength (½) MS medium (pH 5.7) containing 1% sucrose and 0.6% agar, with appropriate antibiotics when necessary. Plated seedlings were grown in a growth chamber at 24/22 °C (day/night) with a 16/8 h light/dark photoperiod. Pant materials created in this study are listed in Table S1.

### Plasmid construction and plant transformation

To generate CRISPR constructs, gRNA spacer sequences and primers were designed using CRISPR-P 2.0 (*55*). The gRNA sequences were subsequently inserted into the high-efficiency CRISPR/Cas9 vector pAGM55273 (*56*) via *Bsa*I-mediated Golden Gate cloning. For the construction of *pUBQ10:LecRK-I.8-DN*, the dominant-negative fragment of *lecrk-I.8-2*, corresponding to amino acids 1-383 of LecRK-I.8, was amplified from Col-0 genomic DNA and inserted into the multiple cloning site (MCS) of the pG20-HYG vector (*57*) using In-Fusion cloning (Takara). To generate *35S:eNahG-GFP*, the *NahG* coding sequence (CDS) was amplified from a plasmid, with a forward primer containing the nucleotides encoding the PR1 signal peptide. The PCR product was digested with *Sac*I and *Xba*I and ligated into the pCAMBIA1300-GFP vector. To generate complementation constructs for the *lecrk-I.8-2* and *lecrk-VI.2-1* mutants, the *LecRK-I.8* and *LecRK-VI.2* CDSs were amplified from Col-0 genomic DNA and cloned into the MCS of pG20-HA vector via In-Fusion cloning. The *pG20-pUBQ10-lecrk-I.8ᴷ³⁵⁷ᴱ-HA*, *pG20-pUBQ10-lecrk-VI.2³^A^-HA*, and *pG20-pUBQ10-lecrk-VI.2^Δloop^-HA* constructs were generated by site-directed mutagenesis using a three-fragment In-Fusion cloning strategy. For *N. benthamiana* transient expression vectors, the extracellular domains of *CARD1*, *BAK1*, *LecRK-I.8*, *LecRK-V.5*, and *LecRK-VI.2*, including their native signal peptide sequences, were amplified from *Arabidopsis* cDNA. The extracellular domains of *lecrk-VI.2^3A^* and *lecrk-VI.2^Δloop^* were amplified from the *pG20-pUBQ10-lecrk-VI.2^3A^-HA*, and *pG20-pUBQ10-lecrk-VI.2^Δloop^-HA* plasmids, respectively. His-tag sequences were added to the reverse primers. The resulting PCR products were inserted into the pHREAC10 vector (*58*), an optimized construct for transient expression in *N. benthamiana*, using In-Fusion cloning. To construct *pUBQ10:cytAEQ*, the *cytAEQ* CDS was amplified from DNA of cytosolic aequorin transgenic plants and inserted into the pG20-HYG vector using In-Fusion cloning. For purification of recombinant NahG protein, *NahG* CDS was amplified from DNA of *NahG* transgenic plants and cloned into the pET28 vector via In-Fusion cloning. To construct *pCAMBIA1300s-AHA2-mCherry*, *AHA2* CDS was amplified from Col-0 cDNA and inserted into pCAMBIA1300s-mCherry vector using In-Fusion cloning. All primers used in this study are listed in Table S2. All constructs were verified by Sanger sequencing or whole plasmid sequencing prior to use. *Arabidopsis* transformation was performed using the floral dip method (*59*). T_1_ transgenic plants were selected on ½ MS medium containing appropriate antibiotics. Lines with single insertion were identified in the T_2_ generation, and homozygous plants were selected in T_3_. All experiments were conducted using seeds from homozygous T_3_ plants unless otherwise specified. CRISPR/Cas9 transgenic lines were genotyped in the T_2_ generation, and homozygous mutants were confirmed by sequencing.

### Bacterial strains and culture conditions

*Escherichia coli* strain XL1-Blue was cultured at 37 °C in LB medium supplemented with appropriate antibiotics for cloning. For *Arabidopsis* transformation, *Agrobacterium tumefaciens* strain GV3101 was used for all constructs except those based on pG20 vectors, for which GV3101(*pSoup*) was used*. A. tumefaciens* strain EHA105 was used for transient expression in *N. benthamiana*. Bacterial pathogen strains were cultured at 28 °C in King’s B medium (2% proteose peptone, 0.15% K_2_HPO_4_, 6 mM MgSO_4_, and 1.5% glycerol) supplemented with appropriate antibiotics and grown overnight to exponential phase. Cultures were then centrifuged at 3,000 × g for 5 min to collect cells. Pellets were resuspended in 1 mM MgCl_2_ and diluted to desired OD_600_ concentrations for leaf inoculation experiments. The *Pseudomonas syringae* pv. *maculicola* ES4326 strain carrying the *luxCDABE* bioluminescence cassette (*Psm_lux*) was previously described (*7*, *60*).

#### SA and NAD^+^ treatments

Sodium salicylate (SA) (Sigma-Aldrich) and NAD^+^ (Research Products International) were dissolved in double-distilled water (ddH_2_O) to the indicated final concentrations. The pH of the NAD^+^ solution was adjusted to 5.7 using NaOH, and the solution was freshly prepared immediately before each experiment. For SA-and NAD^+^-induced local resistance, 0.5 mM SA or 0.2 mM NAD^+^ was infiltrated into two leaves of 4-week-old *Arabidopsis* plants. Water (ddH_2_O) was used as the mock control. Unless otherwise specified, the treated leaves were inoculated with a *Psm* suspension in 1 mM MgCl_2_ (OD_600_ = 0.001) 4 h post-treatment. For NAD^+^-induced systemic resistance, 0.5 mM NAD^+^ was infiltrated into three lower leaves on 4-week-old *Arabidopsis* plants. Water (ddH_2_O) served as the mock control. Four h after treatment, one upper leaf per plant was inoculated with *Psm* (OD_600_ = 0.001). Bacterial populations were quantified and disease symptoms photographed at 2.5 d post inoculation (dpi). For analysis of SA-induced transcriptional reprogramming, 0.5 mM SA or ddH_2_O (mock) was infiltrated into leaves on 4-week-old plants. Four h later, the treated leaves were harvested for RNA extraction, followed by RNA-seq or quantitative PCR (qPCR) analysis.

#### Analysis of basal resistance and local defense gene expression

To assess basal resistance to *Psm*, two leaves per 4-week-old plant were infiltrated with a *Psm* suspension (OD_600_ = 0.0002). Bacterial populations were measured at 0 dpi to confirm equal inoculum levels across samples and at 3 dpi to evaluate pathogen growth in the plants. Disease symptoms were photographed at 3 dpi. For local defense gene expression, four-week-old plants were infiltrated with *Psm* (OD_600_ = 0.001) or 1 mM MgCl_2_ (mock). At 24 h post-infiltration, treated leaves were harvested for RNA extraction.

#### Biological induction of SAR

Three lower leaves on each 4-week-old *Arabidopsis* plant were infiltrated with either 1 mM MgCl_2_ (mock) or *Psm* (OD_600_ = 0.004) to induce SAR. After 48 h, one upper (systemic) leaf per plant was either harvested for RNA extraction or challenged with *Psm* (OD_600_ = 0.001). Bacterial populations were quantified, and disease symptoms were documented by photography at 2.5 dpi.

#### Quantification of pathogen growth

Pathogen growth was quantified using either the traditional colony-forming unit (CFU) method or a bioluminescence-based (relative light unit, RLU) method as previously described (*7*).

#### RNA extraction and qPCR analysis

Total RNA was extracted as previously described (*7*). RNA concentration and quality were assessed using a NanoDrop spectrometer. For reverse transcription (RT), total RNA was treated with the Turbo DNA Free Kit (Invitrogen, Cat#AM1907) at 37 °C for 30 min. DNase was then inactivated, and RT was performed using the LunaScript^®^ RT SuperMix Kit (NEB, #E3010) according to the manufacturer’s instructions. The resulting cDNA was diluted and used for qPCR, which was carried out using SYBR Green PCR Master Mix (Applied Biosystems, Cat#4309155) on a QuantStudio 3 Real-Time PCR system (Applied Biosystems) following the manufacturer’s protocol. Relative gene expression was calculated using the 2^-ΔCt^ method, with *UBQ5* as the internal control. Primers used for qPCR are listed in Table S2.

#### RNA-seq, data analysis and visualization

RNA sequencing was performed at the UF ICBR NextGen Sequencing Core (https://biotech.ufl.edu/next-gen-dna/; RRID:SCR_019152) as previously described (*7*). The genome of *A. thaliana* (version TAIR10) from the database of ENSEMBL was used as the reference sequences for RNA-seq analysis. RNA-seq was conducted with three biological replicates per genotype and treatment. The gene expression levels were analyzed by a DESeq2-based R pipeline (*61*). Hierarchical clustering and heatmaps were calculated and visualized by the Complexheatmap R package (*62*). Gene ontology (GO) enrichment analysis and visualization were conducted using the clusterProfiler (*63*), org.At.tair.db, and ggplot2 R packages. Venn diagrams were generated using the R package VennDiagram.

#### SA measurement

Total (intracellular and extracellular) metabolites were extracted as previously described (*64*). For apoplastic (extracellular) metabolite extraction, leaves were surface-cleaned and vacuum-infiltrated with ddH_2_O using a 60-mL syringe. Residual surface water was carefully removed with Kimwipes. Leaves were then stacked and centrifuged at 500 × *g* for 10 min at 4 °C to collect apoplastic washing fluid (AWF). AWF samples were clarified by centrifugation, supplemented with internal standards (500 ng each), and dried using a ScanSpeed vacuum centrifuge. For intracellular metabolite extraction, after removal of apoplastic fluid as described above, the remaining leaf tissue was frozen in liquid nitrogen, ground to a fine powder, and extracted twice with 1 mL of methanol/ddH_2_O (80:20, v/v). A 600 μL aliquot of the combined extracts was spiked with internal standards (500 ng each), then evaporated to dryness using a ScanSpeed vacuum centrifuge.

Dried samples were subjected to derivatization followed by quantification using gas chromatography-mass spectrometry (GC-MS) as described (*64*) with minor modifications. One microliter of each sample was injected into a Thermo Trace 1610 Gas Chromatograph equipped with a Thermo GC TG-5SilMS column (30 m × 0.25 mm × 0.25 μm). Mass spectra were recorded using an ISQ Single Quadrupole Mass Spectrometer (Thermo Scientific). The quantitative analysis was performed in the Chromeleon 7 software. Specific selected ion chromatograms of analytes and internal standards were integrated as follows: SA (m/z 267) was related to D4-SA (m/z 271).

#### Expression and purification of ectodomain proteins

Apoplastic expression and purification of the extracellular domain proteins from *N. benthamiana* AWF were performed following previously described methods (*65*) with modifications. Briefly, *A. tumefaciens* strains carrying the respective genes were cultured overnight at 28°C in LB media supplemented with 100 μg/mL rifampicin and 50 μg/mL kanamycin. The bacterial cells were harvested by centrifugation at 3,000 × g for 5 min and washed once with agroinfiltration buffer (10 mM MES/NaOH, pH 5.6, 10 mM MgCl_2_, 150 μM acetosyringone). Cells were subsequently resuspended in the agroinfiltration buffer to an optical density (OD_600_) of 0.5 and incubated at room temperature in the dark for 2 h. The resulting suspension was infiltrated into the leaves of 5-week-old *N. benthamiana* plants using a 1 mL needleless syringe. Three d post-infiltration, leaves were harvested and submerged in 150 mL vacuum-infiltration buffer (20 mM Tris/HCl, pH 6.8, 0.01% Tween-20, 1 mM PMSF). Leaves were infiltrated under vacuum in a vacuum desiccator, removed from the buffer, and excess liquid was gently blotted dry using paper towels. The leaves were then rolled, placed in 20-mL syringes (without plungers), and positioned inside 50-mL centrifuge tubes. AWFs were collected by centrifugation at 1,500 × g and 4°C for 10 min and further clarified by additional centrifugation at 10,000 × g and 4°C for 10 min. His-tagged proteins were purified from the clarified AWFs using HiTrap TALON crude columns (Cytiva) according to the manufacturer’s protocol. Protein concentrations were quantified by Bradford assay. Purified proteins were desalted and concentrated using Vivaspin ultrafiltration units (10-kDa molecular weight cut-off). Finally, proteins were aliquoted, snap-frozen in liquid nitrogen, and stored at-80°C until further use. SDS-PAGE analysis was conducted to confirm the purity and molecular weights of the purified proteins.

#### [^3^H]-SA binding assay

[^3^H]-SA binding assays were performed using size exclusion chromatography as previously described with modifications (*14*). Size exclusion columns were prepared by loading 0.13 g of Sephadex G-25 (GE Healthcare) into QIAGEN shredder columns. Columns were pre-equilibrated with SA binding buffer (20 mM MES/KOH pH5.7, 150 mM NaCl, 1 mM CaCl_2_, 1 mM MnCl_2_, 0.1% Tween-20) overnight at 4°C, and excess buffer was removed by centrifugation at 750 × g for 2 min. Binding reactions were performed by combining 2.5 μg of each protein with 200 nM [^3^H]-SA (American Radiolabeled Chemicals; specific activity 30 Ci/mmol), after which SA binding buffer was added up to 25 μL. For competition assays, a 10,000-fold excess of unlabeled SA was added. The reaction mixtures were incubated at room temperature for 30 min. Following incubation, samples were loaded onto the size exclusion columns and immediately centrifuged as described above to separate protein-bound and free [^3^H]-SA. The resulting flow-through was collected. Radioactivity in the eluates was measured using a scintillation counter (LS6500; Beckman Coulter).

#### Microscale thermophoresis assay

Microscale thermophoresis (MST) assays were conducted using a Monolith NT.115 instrument (NanoTemper Technologies). Purified ectodomain proteins were fluorescently labeled using the RED-NHS 2nd Generation labeling kit (NanoTemper Technologies), following the manufacturer’s instructions. Labeled proteins were eluted in MST buffer (20 mM MES/ KOH pH5.7, 150 mM NaCl, 1 mM CaCl_2_, 1 mM MnCl_2_, 0.1% Tween-20). A final concentration of 50 nM labeled proteins with or without 50 nM unlabeled eBAK1-His were mixed with a serial dilution (ranging from 4.6 × 10^-8^ M to 1.5 × 10^-3^ M) of a specific ligand. SA, catechol, and benzoic acid (BA) were prepared in MST buffer, and PC (1,2-dipalmitoleoyl-sn-glycero-3-phosphocholine, PC (16:1/16:1), Avanti Polar Lipids, 850358) was dissolved in 1% DMSO, 0.5% glycerol and 0.1% Tween 20 by sonication (*36*). Binding reactions were incubated at room temperature for 1 h. Samples were then loaded into premium capillaries (NanoTemper Technologies, MO-K025) and analyzed at room temperature using 40% LED power (RED channel) and medium MST power. Data analysis and dissociation constant (*K*d) calculations were performed using MO.Affinity Analysis software (version 2.3).

#### Phos-tag immunoblotting

Phos-tag immunoblotting was performed following a previously described protocol (*66*).

#### Phosphoproteomics

Leaves from 4-week-old *Arabidopsis* plants were vacuum-infiltrated with either 0.5 mM SA or water (mock) using a 50 mL syringe. Treated leaves were incubated in the respective solutions for 15 min and then snap-frozen in liquid nitrogen and stored at-80°C until use. Five hundred μg of total protein from each sample were digested using the filter-aided sample preparation (FASP) method (*67*). Phosphopeptides were enriched using homemade Fe-IMAC tips as previously described (*68*). Dried phosphopeptides were dissolved in 12 µL 0.1% trifluoroacetic acid (TFA), and 8 µl were loaded onto a 50cm C18 capillary column (Thermo PepMap Cat# ES75500PN) with a precolumn on the Thermo Vanquish Neo UHPLC. The peptides are gradually eluted over a 2-h gradient: starting with Solution B (80% ACN/0.1% FA) at 2%, increasing to 4%, 28%, 42%, and 55% over 0.5 min, 90 min, 15 min, and 6 min, respectively, followed by a wash step with 99% of Solution B. The peptides were analyzed on a Thermo Orbitrap Exploris 240 mass spectrometer with the following settings: MS1 was acquired at a resolution of 120k over a scan range of 350-1,600 m/z with an AGC target of 3×10^6^; MS/MS spectrum was collected in a 3-s cycle with the dynamic exclusion duration of 45 s, a resolution of 30k, and a normalized AGC target of 2×10^5^; HCD collision energy was set to 26%. The Thermo raw data files were analyzed using MSFragger in the FragPipe proteomics pipeline (v22.0) with default settings for label free phosphoproteomics except for an updated precursor mass tolerance of-10 to 10 ppm (*69*). The peptides were searched against the *Arabidopsis* database (Araport11) (*70*). Quantification of peptides were conducted using IonQuant (*71*) included in FragPipe with match between runs (MBR) enabled. The MaxLFQ intensity was used to compare between treatment groups. Phosphoproteomic data were filtered to retain phosphosites with at least two valid values in at least one experimental group. Intensities were log_2_ transformed and normalized by using the mode method (subtraction most frequent value). Missing values were imputed using a mean-shift approach (downshift: 1.8; width: 0.3) to simulate the detection limit while minimizing noise. Statistical significance was evaluated using the limma package (*72*) with empirical Bayes moderation. Differentially regulated phosphosites were defined by |log_2_(fold change)| > 0.5 and *P* value < 0.05. Data visualization was performed with ggplot2 and ComplexHeatmap (*62*). GO analysis and visualization were conducted using the clusterProfiler (*63*), org.At.tair.db, and ggplot2 R packages.

#### ROS burst assay

The ROS burst assay was performed as described (*73*). Eight leaf discs (4 mm in diameter) per genotype/treatment were collected from 4 leaves of 4 plants and floated overnight on 100 µL of water in darkness. The next day, water was replaced with ROS assay solution (100 µM luminol (Sigma, A8511) and 20 µg mL^-1^ horseradish peroxidase (Sigma, 77332)) with or without elicitors. Luminescence was immediately recorded as relative light units (RLU) using a GloMax plate reader (Promega).

#### Calcium burst assay

Cytosolic Ca²⁺ bursts were measured using 10-day-old seedlings or leaf discs from 4-week-old plants of Col-0 stably expressing AEQ, or protoplasts transiently expressing AEQ, as previously described (*74, 75*). Briefly, single seedling or leaf disc was transferred into separated wells of white 96-well plates. Each of the wells contained 50 µL of CTZ buffer, including 10 µM coelenterazine (Nanolight technology), 2 mM MES, and 10 mM CaCl_2_ (pH 5.7). The plates were incubated at room temperature overnight in the dark, followed by the addition of 2 mM MES (pH 5.7) as mock or 1 mM SA in MES buffer (pH5.7, 2-fold of the final concentration) directly applied into the wells using a multichannel pipette. RLUs were measured with the Glomax discover luminometer. For Ca²⁺ burst assays in protoplasts, protoplasts were transfected with *pG20:AEQ* and incubated overnight in WI_Ca_ solution (4 mM MES pH 5.7, 0.5 M mannitol, 20 mM KCl, 2 mM CaCl_2_). On the following day, coelenterazine was added to 10 µM final concentration and protoplasts were incubated for one to four hours. Then, 100 µl of protoplast solution containing approximately 2×10^4^ protoplasts were loaded into single wells of 96-well plates and incubated 30 min in the dark before starting the Ca^2+^ measurements. Then 25 µL WI_Ca_ buffer or 1.25 mM SA in WI_Ca_ buffer (pH5.7, 5-fold of the final concentration) was added to the protoplasts and RLUs were recorded with the Glomax discover luminometer. Data are presented as R/R_0_, where R_0_ represents RLU means before the addition of elicitors.

#### MAPK activation assay

For seedling-based assays, 10-day-old seedlings grown on ½ MS medium were transferred to water and incubated overnight in the dark. The following day, seedlings were treated with water or elicitor solutions for the indicated times. For assays using mature plants, leaf discs were collected from 4-week-old soil-grown plants, floated on water overnight, and then syringe-vacuum infiltrated with water or elicitor solutions for the indicated times. Following treatment, seedlings or leaf discs were immediately frozen in liquid nitrogen and ground to a fine powder. Total proteins were extracted by homogenization in ice-cold protein extraction buffer (25 mM HEPES pH 7.4, 150 mM NaCl, 2 mM EDTA, 1 mM sodium fluoride, 1 mM sodium orthovanadate, and protease inhibitor cocktail). Extracts were subjected to SDS-PAGE and immunoblotting with anti-p44/42 MAPK/pERK (Cell Signaling Technology, 9102) to detect phosphorylated MAPKs.

#### Recombinant NahG protein purification and plant treatment

The NahG protein was expressed in *E. coli* strain Rosetta and purified using HisPur™ Cobalt Resin (Thermo Scientific, 89965) according to the manufacturer’s instructions. The purified protein was desalted using a PD-10 column and stored at-80 °C until use. For plant treatments, purified NahG protein was diluted in 10 mM MgCl₂ and infiltrated into 4-week-old plants together with *Psm_lux* (OD_600_ = 0.0002) for the initial treatment. Twelve h later, NahG protein was infiltrated a second time. Plants infiltrated with 10 mM MgCl₂ served as solvent controls.

#### Laser confocal microscopy

AHA2-mCherry and eNahG-GFP were transiently co-expressed in *N. benthamiana* for 3 days as described (*76*). Fluorescence images were acquired using a Leica Stellaris confocal laser scanning microscope with excitation and detection settings optimized for GFP and mCherry.

#### *In vivo* co-immunoprecipitation assay

Transgenic plants were treated with SA or water (mock) for 10 min. Treated leaves were immediately frozen in liquid nitrogen and ground into a fine powder. The tissue powder was homogenized in co-IP buffer (25 mM HEPES pH 7.4, 150 mM NaCl, 0.5% Triton X-100, 1 mM sodium fluoride, 1 mM sodium orthovanadate, and protease inhibitor cocktail) and clarified by centrifugation at 4 °C. Cleared lysates were incubated with HA-Trap (ChromoTek) for 2 h at 4 °C, followed by five washes with wash buffer (25 mM HEPES pH 7.4, 150 mM NaCl, 0.1% Triton X-100). The washed beads were resuspended in 2× SDS loading buffer and denatured for subsequent SDS-PAGE and immunoblot analysis. BAK1 was detected using an anti-BAK1 antibody (Agrisera, AS121858).

#### Computational modeling of eLecRK-VI.2-SA binding

Structural models of the LecRK-VI.2 extracellular domain were generated using Boltz-2 (*77*) by providing the protein sequence together with the SMILES representation of SA. Complexes were generated using default parameters, and candidate ligand-binding modes were selected based on proximity to the receptor surface and the presence of stabilizing interactions. Structural models were similarly generated for the F111A/R165A/R177A triple mutant and a V155-I166 loop-deletion variant in the absence of SA. Protein-ligand complexes were subsequently prepared and subjected to energy minimization using standard protocols in AMBER (*78*) to relieve steric clashes and optimize geometry. Predicted ligand-binding residues and structural features were evaluated experimentally by mutagenesis and biochemical binding assays. Full details of model generation, ligand preparation, and minimization are provided in the Supplementary Text.

#### Measurement of stomatal aperture

Stomatal aperture measurements were performed on epidermal peels taken from the abaxial side of leaves of 4-week-old plants, following previously described methods with slight modifications (*79*). Three epidermal peels from three individual plants were incubated in stomatal opening buffer containing 10 mM KCl and 25 mM MES-KOH (pH 6.15), under light in the growth room for 3 h to induce maximum stomatal opening. To evaluate stomatal closure, peels were treated SA (0.5 mM) or ABA (5 μM). Corresponding mock treatments included water for SA and 0.1% DMSO for ABA. Stomatal apertures were measured 1 h after treatment with a microscope. The width and length of each stomatal pore were measured using ImageJ, and the stomatal aperture index was calculated as the ratio of width to length.

## Statistical analysis

Sample sizes were not predetermined using statistical methods, and no blinding or randomization was applied. Quantitative data are presented as mean ± SEM or as box plots, where boxes indicate the interquartile range, whiskers represent the minimum and maximum values, and the median is shown as a solid line. Statistical significance was assessed using two-sided Student’s t-test, or one-way or two-way ANOVA followed by Tukey’s post hoc test, conducted in R. The number of biologically independent replicates is provided in the figure legends, and *P* and false discovery rate-adjusted *P* values are reported. The disease resistance results presented in this study are derived from one experimental dataset. These results were confirmed in at least three independent experiments conducted at different times. The gene expression results were from analyses of three independent biological samples. Western blotting, SA quantification, and stomatal closure assays were independently repeated three times with similar results.

## Acknowledgments

We thank Dr. Cyril Zipfel at University of Zurich and Dr. Ping He at University of Michigan for providing *bak1-5 bkk1* seeds, Dr. Marc Knight at Durham University and Dr. Gary Stacey at University of Missouri for providing ctyAEQ seeds, and Dr. Xin Li at University of British Columbia for sharing with us *35S:nahG* seeds. The T-DNA insertion lines SAIL_1145_B10 (*fin4-3*), SALK_005125 (*lecrk-I.8-2*), SALK_070801 (*lecrk-VI.2-1*), SAIL_1146_B02 (*lecrk-VI.2-2*), SAIL_796_E11 (*lecrk-VI.2-3*), SALK_026551 (*svp-31*), and SALK_072930 (*svp-32*) were ordered from the Arabidopsis Biological Resource Center.

## Author contributions

Conceptualization: QL, ZM

Methodology: QL, CL, XW, ZM

Investigation: QL, MZ, CS, CRL, BAM, FY, YZ, XW

Visualization: QL, ZM

Funding acquisition: ZM, MZ

Project administration: ZM

Supervision: ZM

Writing – original draft: QL, ZM

Writing – review & editing: QL, ZM

## Data, code, and materials availability

The authors declare that all data supporting the findings of this study are available within the manuscript and its supplementary files or are available from the corresponding author upon request. RNA-seq data generated as part of this study have been deposited in the Gene Expression Omnibus repository under accession codes GSE295980, GSE296271, GSE312725, and GSE314895. The mass spectrometry proteomics data have been deposited in the MassIVE repository (dataset identifier: MSV000100124) and the ProteomeXchange Consortium (dataset identifier: PXD071648) (*51*).

## Supplementary Materials

Materials and Methods

Supplementary Text

Figs. S1 to S10

Tables S1 to S2

References (*51*-*97*)

Data S1 to S4

