## Supplemental materials for "Extracellular salicylic acid activates immune signaling through cell-surface receptors"

**The PDF file includes:**

Supplementary Text
Figs. S1 to S10
Tables S1 to S2

**Other Supplementary Materials for this manuscript include the following:**

Data S1 to S4

### Supplementary Text

Computational modeling of eLecRK-VI.2-SA binding

**Structural Model Generation**. To construct the structural model, we provided Boltz-2 (*77*) with the amino acid sequence of the extracellular domain of LecRK-VI.2 (HVRAQRTTTNFAFRGFNGNQSKIRIEGAAMIKPDGLLRLTDRKSNVTGTAFYHKPVRLL NRNSTNVTIRSFSTSFVFVIIPSSSSNKGFGFTFTLSPTPYRLNAGSAQYLGVFNKENNGDP RNHVFAVEFDTVQGSRDDNTDRIGNDIGLNYNSRTSDLQEPVVYYNNDDHNKKEDFQL ESGNPIQALLEYDGATQMLNVTVYPARLGFKPTKPLISQHVPKLLEIVQEEMYVGFTAST GKGQSSAHYVMGWSFSSGGERPIADVLILSELPPPPPNKAKKEGLNSQV), together with the SMILES representation of salicylic acid (SA; **O=C([O-])c1ccccc1O)**. Complexes were generated using default Boltz-2 parameters, yielding 100 structural samples. These samples were screened to identify plausible ligand-binding modes based on spatial proximity to the receptor surface and the presence of extensive stabilizing interactions such as hydrophobic and/or hydrogen-bonding interactions. Using the same protocol, we also generated structural models for a Phe111A/Arg165A/Arg177A triple mutant and for a Val155-Ile166 loop deletion variant without SA bound.

**Model and Ligand Preparation.** SA underwent geometry optimization at the B3LYP (*80-83*)/6-31G* (*84-88*) level of theory using Gaussian16 (*89*). Electrostatic potential (ESP) charges were computed at the same level, and restrained electrostatic potential (RESP) charges were subsequently derived using the RESP procedure (*90*) as implemented in the Antechamber module of AmberTools (*78*). A physiologically relevant SA conformer was generated with Schrödinger Maestro's LigPrep (*91*) to assign the appropriate protonation state. Complex topology and coordinate files were constructed using tleap.

**Model Minimization.** Energy minimizations were performed using the CPU implementation of pmemd (*92-94*) in AMBER24 (*78, 95*), with the ff19SB (*96*) protein forcefield. The SA-bound system was solvated in a truncated octahedron containing OPC (*97*) water with a 12.0 Å buffer. Sodium and chloride ions were added to neutralize the system and achieve a physiological salt concentration of 150 mM NaCl. Minimization proceeded through a five-stage protocol designed to sequentially relax solvent, sidechains, and backbone atomic clashes. First, the system underwent an energy minimization of 10 steps of steepest descent followed by 4,990 steps of conjugate gradient, during which all atoms but water molecules were restrained with a force constant of 100.0 kcal/mol/Å$^2$. Next, the system underwent an energy minimization of 10 steps of steepest descent followed by 4,990 steps of conjugate gradient, during which all atoms but water molecules and ions were restrained with a force constant of 100.0 kcal/mol/Å$^2$. Next, the system underwent an energy minimization of 10 steps of steepest descent followed by 4,990 steps of conjugate gradient, during which only the heavy atoms of the protein backbone and ligand were restrained with a force constant of 100.0 kcal/mol/Å$^2$. Next, the system underwent an energy minimization of 10 steps of steepest descent followed by 4,990 steps of conjugate gradient, during which only the heavy atoms of the protein backbone were restrained with a force constant of 100.0 kcal/mol/Å$^2$. Finally, the system underwent an energy minimization of 10 steps of steepest descent followed by 9,990 steps of conjugate gradient, with no restraints present. The resulting minimized structure was used for subsequent analyses.

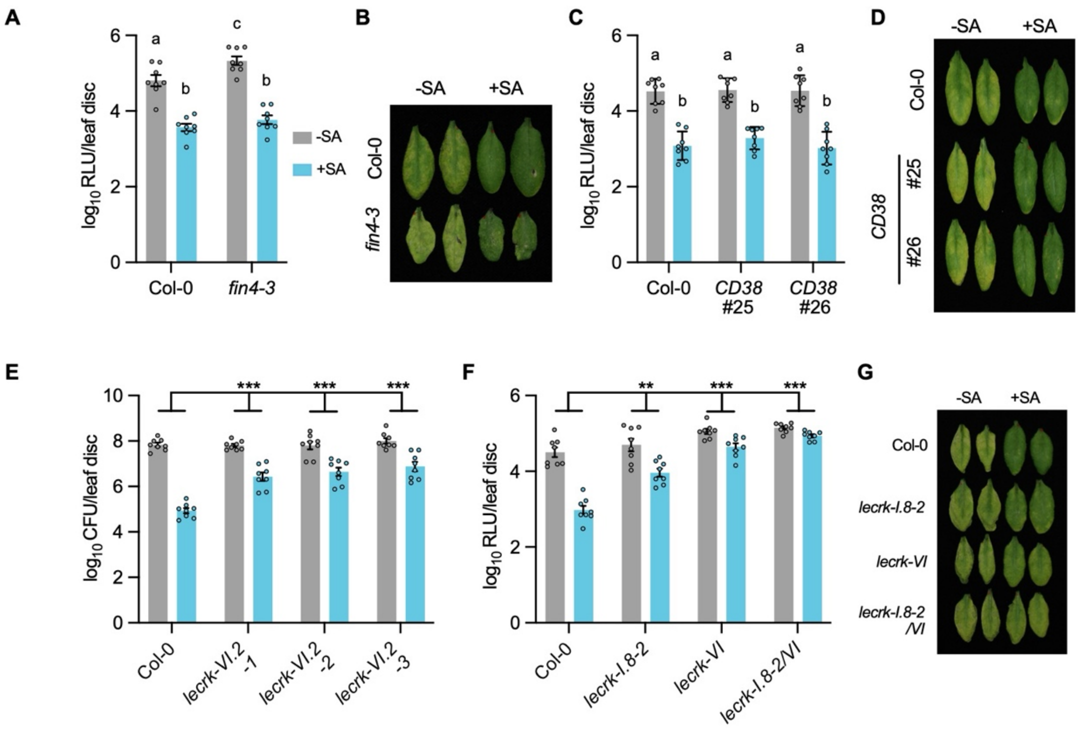


**Fig. S1.**

**SA induces eNAD(P)-independent but LecRK-dependent resistance** (A to D) SA-induced resistance in *fin4-3* and *35S:CD38* (*CD38*) plants. Leaves on 4-week-old Col-0, *fin4-3* (A and B), and *CD38* (C and D) plants were infiltrated with water (-SA) or 0.5 mM SA (+SA). Four h later, the treated leaves were inoculated with *Psm_lux* ($OD_{600}$ = 0.001). Bacterial populations ( A and C) were quantified, and disease symptoms (B and D) were photographed at 2.5 d post-inoculation (dpi). RLU, relative light unit. Bars in (A and C) represent means ± standard error of the mean (SEM; n = 8). Different letters denote significant differences ($P < 0.05$; one-way ANOVA followed by Tukey's test). (E to G) SA-induced resistance in *lecrk* mutants. Plants were treated as in (A). Four h later, the treated leaves were inoculated with non-luminescent *Psm* ($OD_{600}$ = 0.001) (E) or bioluminescent *Psm_lux* ($OD_{600}$ = 0.001) (F). Bacterial populations were quantified by traditional colony-forming unit (CFU) counts (E) or by a bioluminescent assay (F). Disease symptoms (G) were photographed at 2.5 dpi. Bars represent means ± SEM (n = 8). Asterisks denote significant differences (****P* < 0.001, ***P* < 0.01; two-way ANOVA).

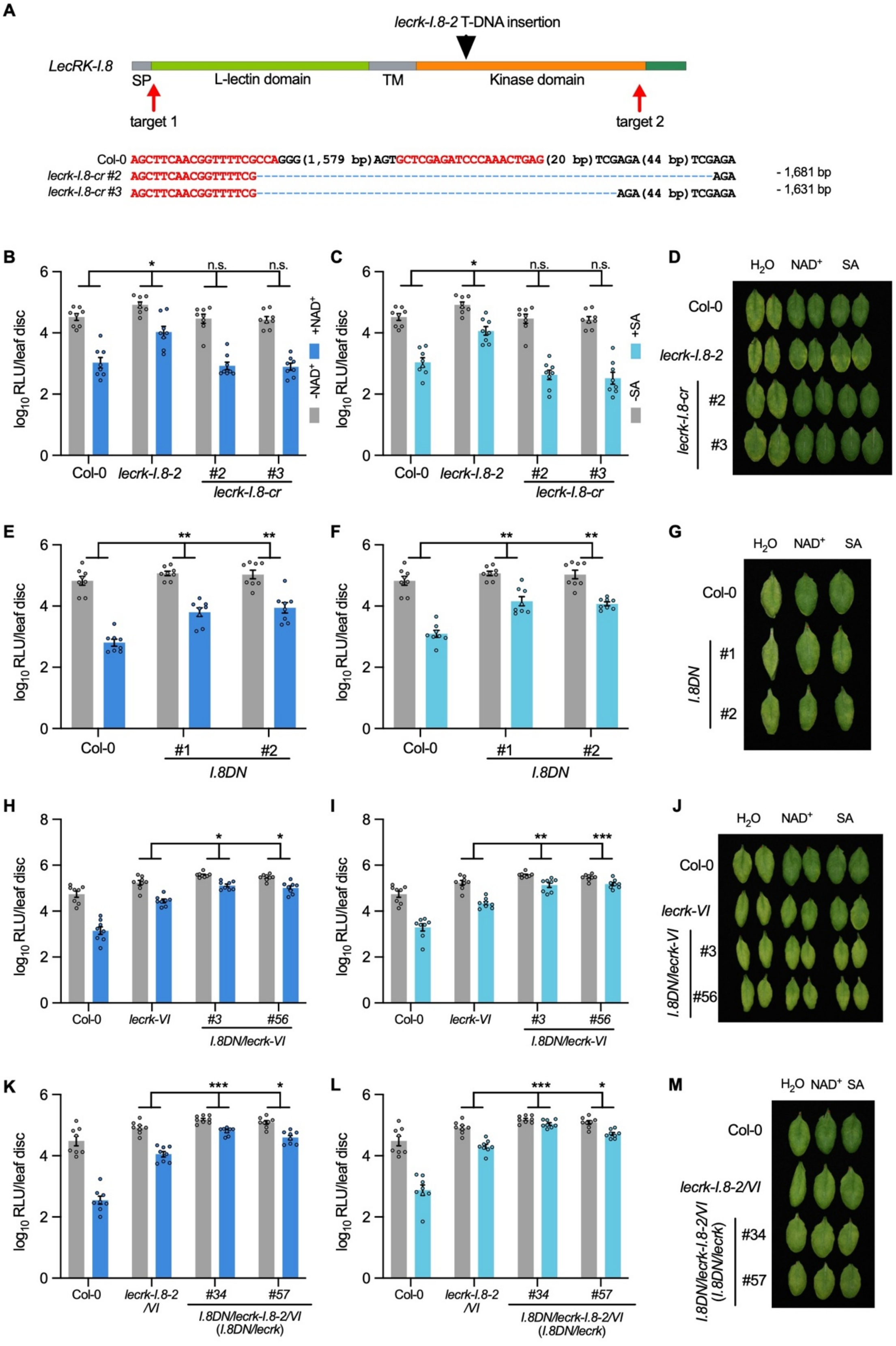
A
lecrk-I.8-2 T-DNA insertion
LecRK-I.8
SP
L-lectin domain
TM
Kinase domain
target 1
target 2
Col-0 AGCTTCAACGGTTTTCGCCAGGG(1,579 bp)AGTGCTCGAGATCCCAAACTGAG(20 bp)TCGAGA(44 bp)TCGAGA
lecrk-I.8-cr #2 AGCTTCAACGGTTTTCG AGA - 1,681 bp
lecrk-I.8-cr #3 AGCTTCAACGGTTTTCG AGA(44 bp)TCGAGA - 1,631 bp
B
C
D
E
F
G
H
I
J
K
L
M
log10 RLU/leaf disc
+NAD+
-NAD+
+SA
-SA
n.s.
Col-0
lecrk-I.8-2
lecrk-I.8-cr
I.8DN
lecrk-VI
I.8DN/lecrk-VI
lecrk-I.8-2/VI
I.8DN/lecrk-I.8-2/VI (I.8DN/lecrk)
H2O
NAD+
SA

**Fig. S2.**

**The T-DNA insertion line *lecrk-I.8-2* is a dominant-negative mutant** (A) Schematic diagram illustrating the T-DNA insertion position in *lecrk-I.8-2* and the CRISPR/Cas9-induced deletions in *lecrk-I.8-cr* (CRISPR) mutants. The black arrowhead denotes the T-DNA insertion position. Red arrows mark the CRISPR gRNA target sites, with the gRNA spacer sequences highlighted in red. Blue dashed lines indicate the deleted regions. Bracketed numbers represent the number of nucleotides omitted in the Col-0 reference sequence, and the number of nucleotides deleted (-) in each *lecrk-I.8-cr* mutant is indicated on the right. (B to D) $NAD^+$- and SA-induced resistance in the *lecrk-I.8-cr* mutants. (E to M) $NAD^+$- and SA-induced resistance in *pUBQ10:lecrk-I.8-2DN* (*I.8DN*) lines in Col-0 (E to G), *lecrk-VI* (H to J), and *lecrk-I.8-2/VI* (K to M) backgrounds. Leaves on 4-week-old plants were infiltrated with water, 0.2 mM $NAD^+$, or 0.5 mM SA. Four h later, the treated leaves were inoculated with *Psm_lux* ($OD_{600}$ = 0.001). Bacterial populations were quantified, and disease symptoms were photographed at 2.5 dpi. Bars represent means ± SEM (n = 8). Asterisks denote significant differences (****P* < 0.001, ***P* < 0.01, **P* < 0.05, n.s., not significant; two-way ANOVA).

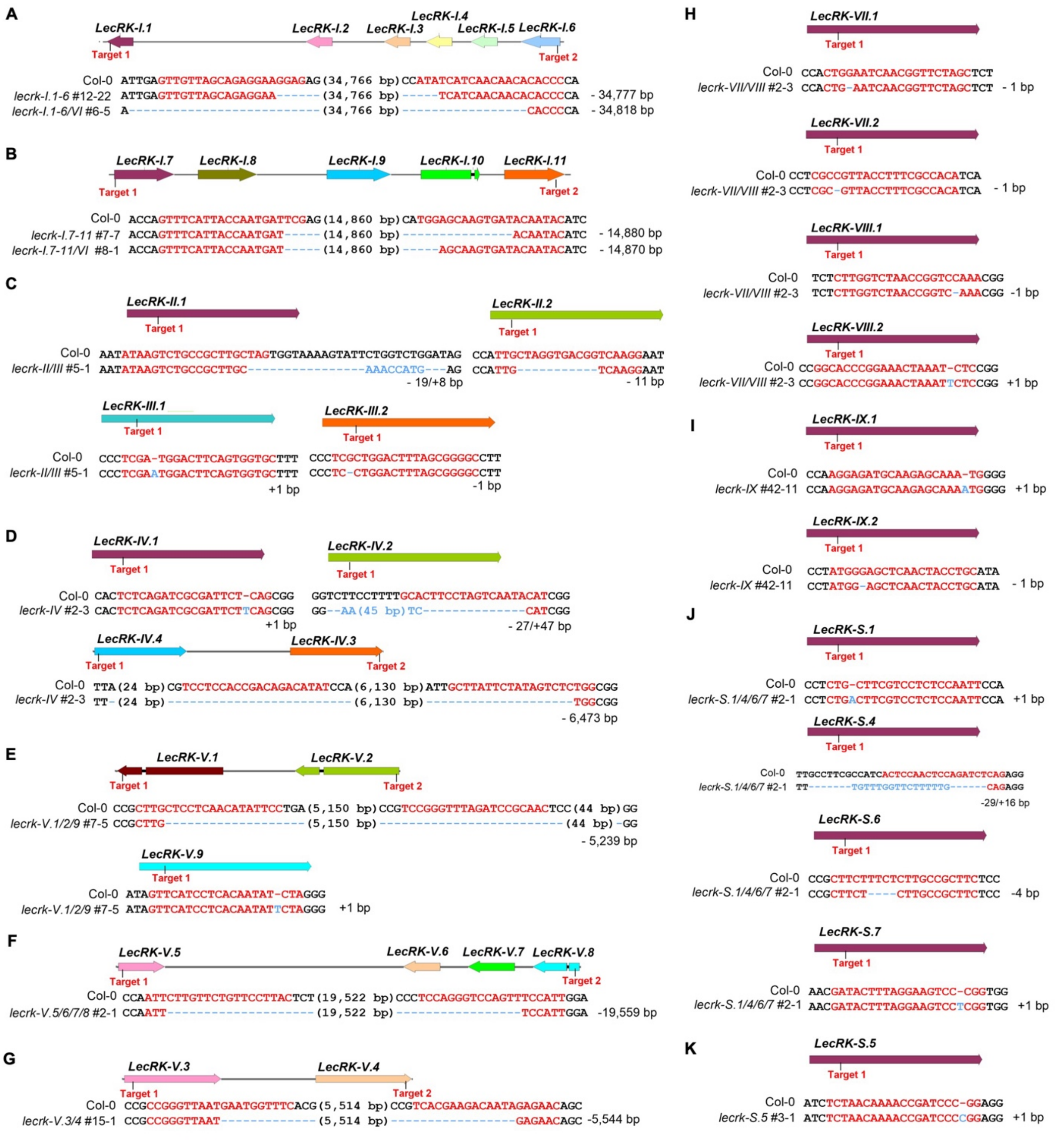


**Fig. S3.**

**Schematic diagrams of CRISPR/Cas9-generated higher-order *lecrk* mutants.** The 43 *Arabidopsis* L-type *LecRK* genes were grouped based on their phylogenetic relationships and genomic clustering, and each group was independently knocked out using CRISPR/Cas9. The groups are defined as follows: (A) *LecRK-I.1-6*, (B) *LecRK-I.7-11*, (C) *LecRK-II/III*, (D) *LecRK-IV*, (E)*LecRK-V.1/2/9*, (F) *LecRK-V.5/6/7/8*, (G) *LecRK-V.3/4*, (H) *LecRK-VII/VIII*, (I) *LecRK-*

*IX*, (J) *LecRK-S.1/4/6/7*, and (K) *LecRK-S.5*. The gRNA target sites are marked, with the gRNA spacer sequences highlighted in red. Blue dashed lines indicate the deleted regions and blue letters are inserted nucleotides. Bracketed numbers represent the number of nucleotides omitted in the Col-0 reference sequence, and the number of nucleotides deleted (-) and/or inserted (+) in each CRISPR mutant is indicated alongside the corresponding sequence.

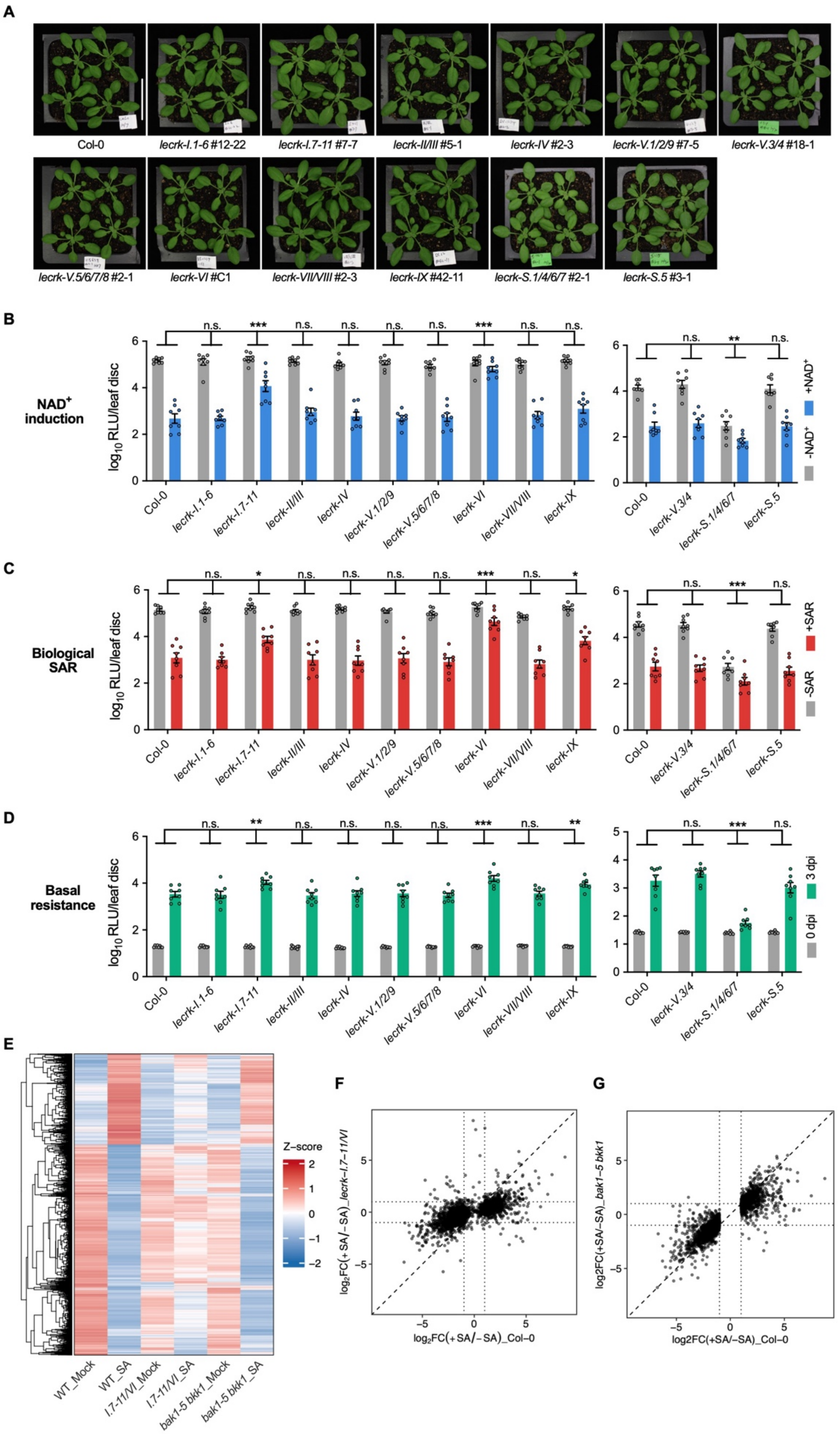
A
Col-0
lecrk-I.1-6 #12-22
lecrk-I.7-11 #7-7
lecrk-II/III #5-1
lecrk-IV #2-3
lecrk-V.1/2/9 #7-5
lecrk-V.3/4 #18-1
lecrk-V.5/6/7/8 #2-1
lecrk-VI #C1
lecrk-VII/VIII #2-3
lecrk-IX #42-11
lecrk-S.1/4/6/7 #2-1
lecrk-S.5 #3-1
B
NAD+ induction
log10 RLU/leaf disc
+NAD+
-NAD+
C
Biological SAR
+SAR
-SAR
D
Basal resistance
3 dpi
0 dpi
E
Z-score
WT_Mock
WT_SA
I.7-11/VI_Mock
I.7-11/VI_SA
bak1-5 bkk1_Mock
bak1-5 bkk1_SA
F
log2FC(+SA/−SA)_lecrk-I.7-11/VI
log2FC(+SA/−SA)_Col-0
G
log2FC(+SA/−SA)_bak1-5 bkk1
log2FC(+SA/−SA)_Col-0

**Fig. S4.**

**Morphology, basal resistance, and induced resistance in higher-order *lecrk* CRISPR mutants, and SA-induced transcriptome changes in *lecrk-I.7-11/VI* and *bak1-5 bkk1*** (A) Morphology of 4-week-old higher-order *lecrk* CRISPR mutants. Scale bar, 4 cm. (B) $NAD^+$-induced resistance to *Psm* in the higher-order *lecrk* mutants. Leaves on 4-week-old plants were infiltrated with water ($-NAD^+$) or 0.2 mM $NAD^+$ ($+NAD^+$). Four h later, the treated leaves were inoculated with *Psm_lux* ($OD_{600}$ = 0.001). Bacterial populations were quantified at 2.5 dpi. (C) Biological induction of SAR in the higher-order *lecrk* mutants. Three lower leaves on each 4-week-old plant were infiltrated with either 1 mM $MgCl_2$ (-SAR) or *Psm* ($OD_{600}$ = 0.004, +SAR) to induce SAR. Two d later, one systemic leaf on each plant was inoculated with *Psm_lux* ($OD_{600}$ = 0.001). Bacterial populations in the systemic leaves were measured at 2.5 dpi. (D) Basal resistance of the higher-order *lecrk* mutants to *Psm*. Leaves on 4-week-old plants were inoculated with *Psm_lux* ($OD_{600}$ = 0.0002), and bacterial populations were quantified at 0 and 3 dpi. Bars represent means ± SEM (n = 8). Asterisks denote significant differences (***$P < 0.001$, **$P < 0.01$, *$P < 0.05$, n.s., not significant; two-way ANOVA). (E to G) SA-induced transcriptional reprogramming in *lecrk-I.7-11/VI* (*I.7-11/VI*) and *bak1-5 bkk1*. The indicated plants were treated with water (mock, -SA) or 0.4 mM SA. Four h later, the treated leaf tissues were collected for RNA-seq analysis. Differentially expressed genes (DEGs) in all genotypes (adjusted *P* value (*P*_adj) < 0.05; $log_2$FC(+SA/-SA) > 1) were extracted and visualized by heatmap (E) and scatter plots (F and G) as described in Methods. FC, fold change.

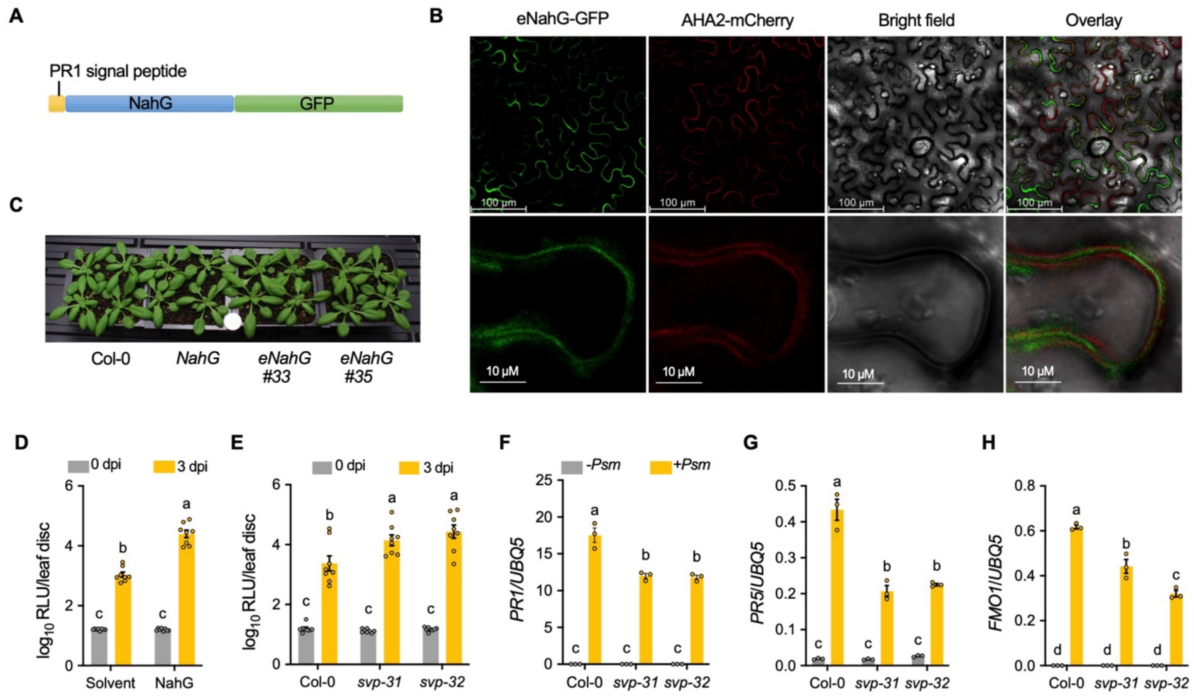


**Fig. S5.**
**Modulation of plant immunity by apoplastic NahG and *svp* mutations** (A) Schematic diagram of the PR1 signal peptide-NahG-GFP fusion protein. (B) Confocal microscopy images showing that AHA2-mCherry and eNahG-GFP are localized in the plasma membrane and the apoplast, respectively, in *N. benthamiana*. (C) Morphology of 4-week-old Col-0, *NahG*, and *eNahG* plants. A U.S. dime was included for scale. (D) Apoplastic infiltration of recombinant NahG protein induces susceptibility. Recombinant NahG (10 µg/mL) was co-infiltrated with *Psm_lux* ($OD_{600}$ = 0.0002) and reinfiltrated at 12 hpi, with 10 mM $MgCl_2$ as the solvent control. (E) Basal resistance of *svp-31* and *svp-32* to *Psm*. Leaves on 4-week-old plants were inoculated with *Psm_lux* ($OD_{600}$ = 0.0002). Bacterial populations were quantified at 0 and 3 dpi. Bars represent means ± SEM (n = 8). (F to H) *Psm*-induced *PR1* (F), *PR5* (G), and *FMO1* (H) expression in *svp-31* and *svp-32*. Four-week-old plants were infiltrated with 1 mM $MgCl_2$ (-*Psm*) or *Psm* ($OD_{600}$ = 0.001). Leaf tissues were collected 24 h later for qPCR analysis. Bars represent means ± SEM (n = 3). Different letters in denote significant differences ($P < 0.05$; one-way ANOVA followed by Tukey's test).

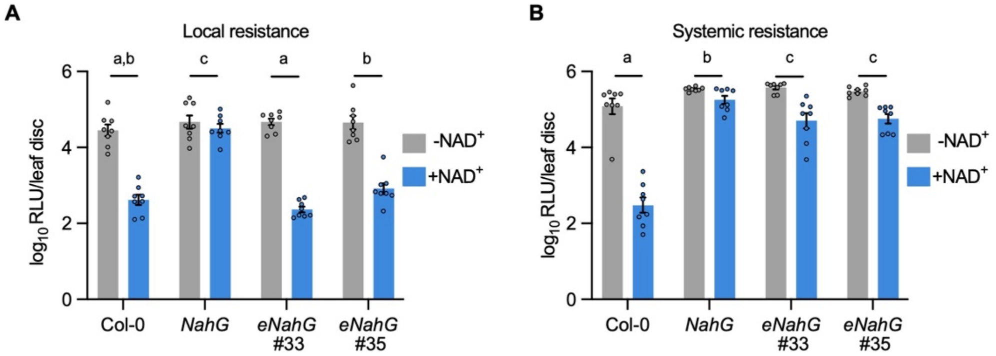


**Fig. S6.**
**NAD$^+$-induced local and systemic immunity in *NahG* and *eNahG* plants** (A) NAD$^+$-induced local resistance. Leaves on 4-week-old plants of the indicated genotypes were infiltrated with water or 0.2 mM NAD$^+$. Four h later, the treated leaves were inoculated with *Psm_lux* ($OD_{600}$ = 0.001). (B) NAD$^+$-induced systemic resistance. Three lower leaves on each 4-week-old plant were infiltrated with water or 0.5 mM NAD$^+$. Four h later, one upper, untreated leaf on each plant was inoculated with *Psm_lux* ($OD_{600}$ = 0.001). Bacterial populations were quantified at 2.5 dpi. Bars represent means ± SEM (n = 8). Different letters denote significant differences ($P$ < 0.05; two-way ANOVA).

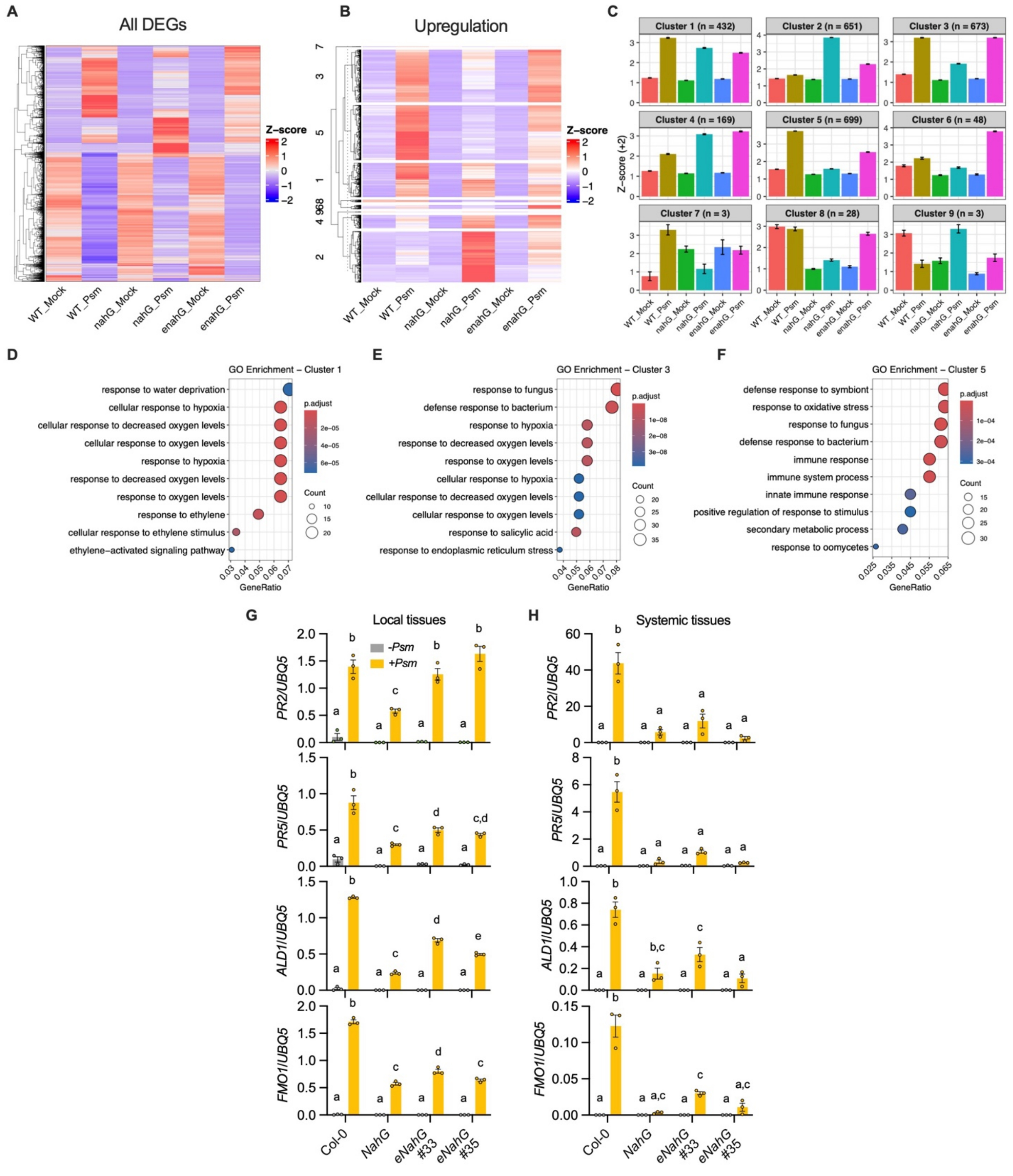


**Fig. S7.**

**Transcriptional changes in *NahG*, *eNahG*, and *lecrk* mutant plants** (A) Heatmap showing DEGs in Col-0, *NahG*, and *eNahG* plants infiltrated/treated with 1 mM $MgCl_2$ (mock) or *Psm* ($OD_{600}$ = 0.001) for 24 h. DEGs were defined as genes with *P*_adj < 0.05 and $|log_2FC| > 2$ relative to mock treatment. (B) Split heatmap showing genes significantly upregulated in *Psm*-infected plants compared to mock-treated controls in Col-0, *NahG*, and *eNahG* ($log_2FC > 2$;

$P$_adj < 0.05). Genes were grouped by hierarchical clustering, and cluster identities are indicated on the left. Gene expression levels (FPKM) in (A and B) were derived from RNA-seq data and normalized by z-score transformation. Rows represent individual genes, and columns represent different genotypes and treatments. The color scale indicates relative expression levels (z-scores). (C) Expression trends of each cluster identified in (B). Bars represent the means ± SEM of normalized expression levels (z-score + 2) for each cluster under each condition. The number of genes in each cluster is indicated. (D to F) Gene Ontology (GO) enrichment analysis of cluster 1 (D), cluster 3 (E), and cluster 5 (F) identified in (B). The top 10 most significantly enriched biological process terms are shown. (G and H) *Psm*-induced defense gene expression in local (G) and systemic (H) tissues of Col-0, *NahG*, and *eNahG* plants. For local tissues, leaves on 4-week-old plants were infiltrated with 1 mM $MgCl_2$ (-*Psm*) or *Psm* ($OD_{600}$ = 0.001) (+*Psm*), and the infiltrated leaves were collected 24 h later. For systemic tissues, leaves on 4-week-old plants were infiltrated with *Psm* ($OD_{600}$ = 0.004) or 1 mM $MgCl_2$, and the upper, untreated leaves were collected 48 h later. Gene expression was analyzed by qPCR. Bars represent means ± SEM (n = 3). Different letters in denote significant differences ($P < 0.05$; one-way ANOVA followed by Tukey's test).

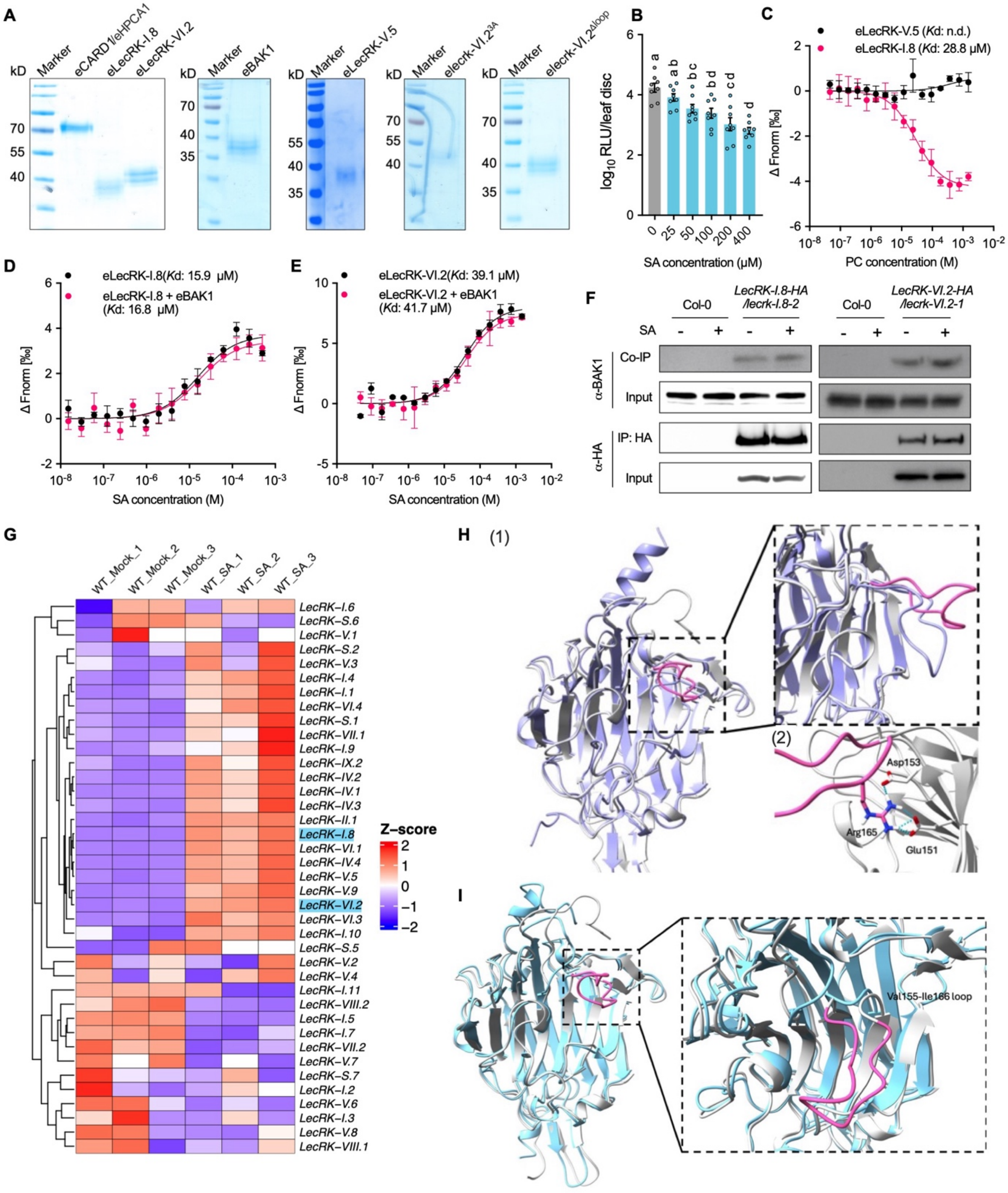


**Fig. S8.**

**Biochemical, molecular, and computational analyses of LecRKs** (A) Coomassie brilliant blue staining of recombinant extracellular domain proteins purified from *N. benthamiana* for MST assays. (B) Dose-dependent induction of *Psm* resistance by SA. Four-week-old Col-0 plants were infiltrated with a concentration series of SA, with water (0 μM) as the control. After 4 h, leaves were challenge-inoculated with *Psm_lux* ($OD_{600}$ = 0.001). Pathogen growth was assessed at 2.5 dpi. Bars represent means ± SEM (n = 8). Different letters denote significant differences ($P$ <

0.05; one-way ANOVA followed by Tukey's test). (C) MST analysis of PC binding to eLecRK-I.8 or eLecRK-V.5. ΔFnorm represents the normalized fluorescence changes of labeled proteins. Data represent means ± SEM (n = 3 independent experiments). $K_d$ values were determined by nonlinear regression, and best-fit values are shown. n.d., not detected. (D and E) The effect of eBAK1 on SA binding affinities of eLecRK-I.8 (D) and eLecRK-VI.2 (E). (F) Co-immunoprecipitation (co-IP) assays examining the effect of SA treatment on interactions between BAK1 and LecRK-I.8 or LecRK-VI.2. Leaves on 4-week-old plants were infiltrated with water (-) or 0.5 mM SA for 10 min, and leaf tissues were subjected to co-IP as described in Methods. (G) Heatmap showing expression of *LecRK* genes in wild-type (WT) Col-0 treated with water (mock) or 0.5 mM SA for 4 h. Three biological replicates per treatment are shown. Gene expression values were derived from RNA-seq data in Fig. 1F and normalized using z-scores. The color scale represents relative expression levels (z-scores). (H) Overlay of predicted extracellular domain structures of wild-type LecRK-VI.2 (gray) and lecrk-VI.$2^{3A}$ (purple) (1). Close-up view of the loop region in wild-type LecRK-VI.2 (pink) and important interactions to maintain SA-binding pocket (2). (I) Overlay of predicted extracellular domain structures of wild-type LecRK-VI.2 (gray) and lecrk-VI.$2^{\Delta loop}$ (blue).

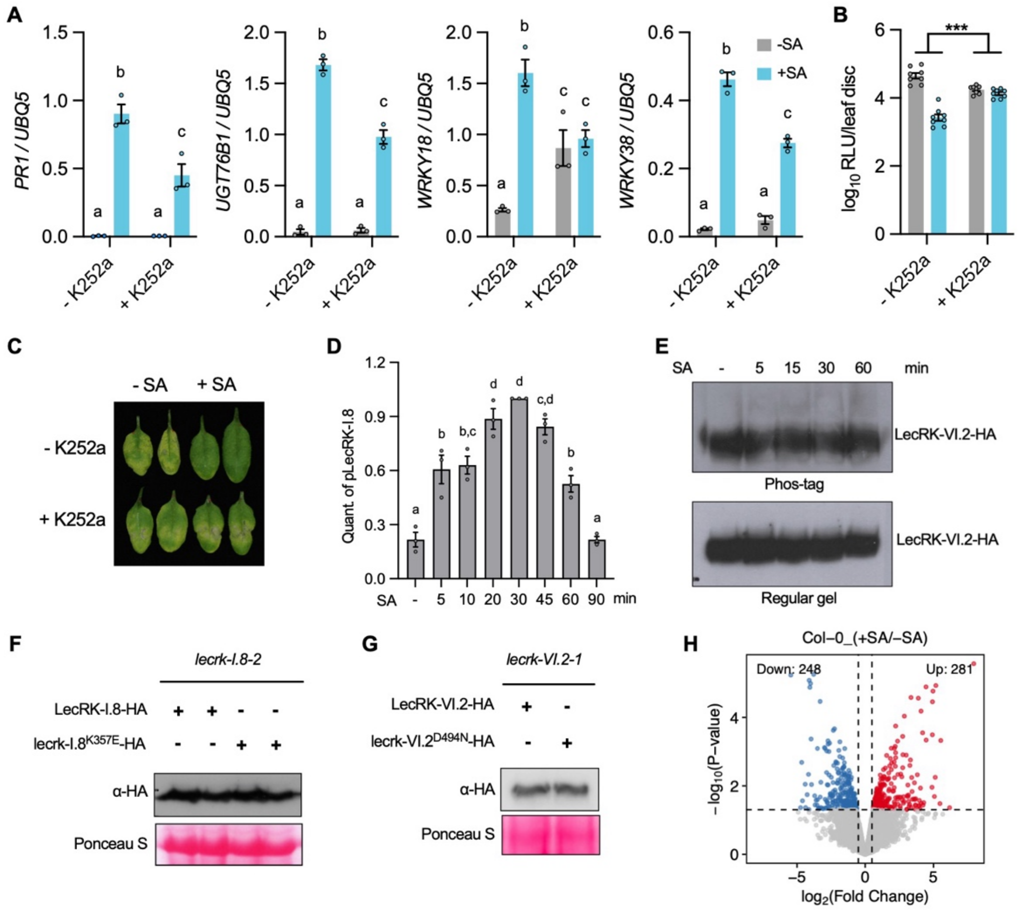


**Fig. S9.**

**SA induces immune responses and protein phosphorylation changes in a kinase activity-dependent manner** (A) The effect of K252a treatment on SA-induced expression of defense-related genes. Leaves on 4-week-old Col-0 plants were infiltrated with water (-SA) or 0.5 mM SA (+SA), either with 0.1% DMSO (- K252a) or 8 µM K252a. The treated leaves were harvested 4 h later for qPCR analysis. Bars represent means ± SEM (n = 3). Different letters denote significant differences (*P* < 0.05; one-way ANOVA followed by Tukey's test). (B and C) The effect of K252a treatment on SA-induced resistance. Leaves on 4-week-old Col-0 plants were infiltrated with water or 0.5 mM SA, either with 0.1% DMSO (- K252a) or 8 µM K252a. Four h later, the treated leaves were inoculated with *Psm* ($OD_{600}$ = 0.001). Bacterial populations (B) were quantified, and disease symptoms (C) were photographed at 2.5 dpi. Bars represent means ± SEM (n = 8). Asterisks denote significant differences (****P* < 0.001; two-way ANOVA). (D) Quantification of pLecRK-I.8 in (Fig. 4B) was performed using ImageJ and is shown as the ratio of the pLecRK-I.8-HA signal in Phos-tag gel to the LecRK-I.8-HA signal in regular SDS-PAGE. The average ratio at 30 min was set as the reference (1) for normalization. Bars represent means ± SEM (n = 3). Different letters indicate significant differences (p < 0.05; one-way ANOVA). (E) Phos-tag assay of LecRK-VI.2-HA following SA treatment. Leaves on 4-week-old *pLecRKVI.2:LecRK-VI.2-HA/lecrk-VI.2-1* plants were vacuum-infiltrated with water (-) or 0.6 mM SA. Total proteins were extracted from leaf tissues collected at the indicated time points (the water-treated sample was collected at 20 min) and separated by $Mn^{2+}$-Phos-tag or regular SDS-PAGE, followed by immunoblotting with anti-HA antibody. (F and G) Protein levels of LecRK-

I.8-HA (F) and LecRK-VI.2-HA (G), as well as their respective kinase-dead variants, in transgenic *lecrk-I.8-2* or *lecrk-VI.2-1* plants expressing the corresponding transgenes. Total proteins were analyzed by immunoblotting with anti-HA antibody. Ponceau S staining of Rubisco was used as the loading control. (H) Volcano plot showing SA-induced rapid and global protein phosphorylation changes in Col-0 revealed by phosphoproteomic analysis. Significantly regulated phosphosites were defined as those with $P < 0.05$ and $|\log_2 FC| > 0.5$.

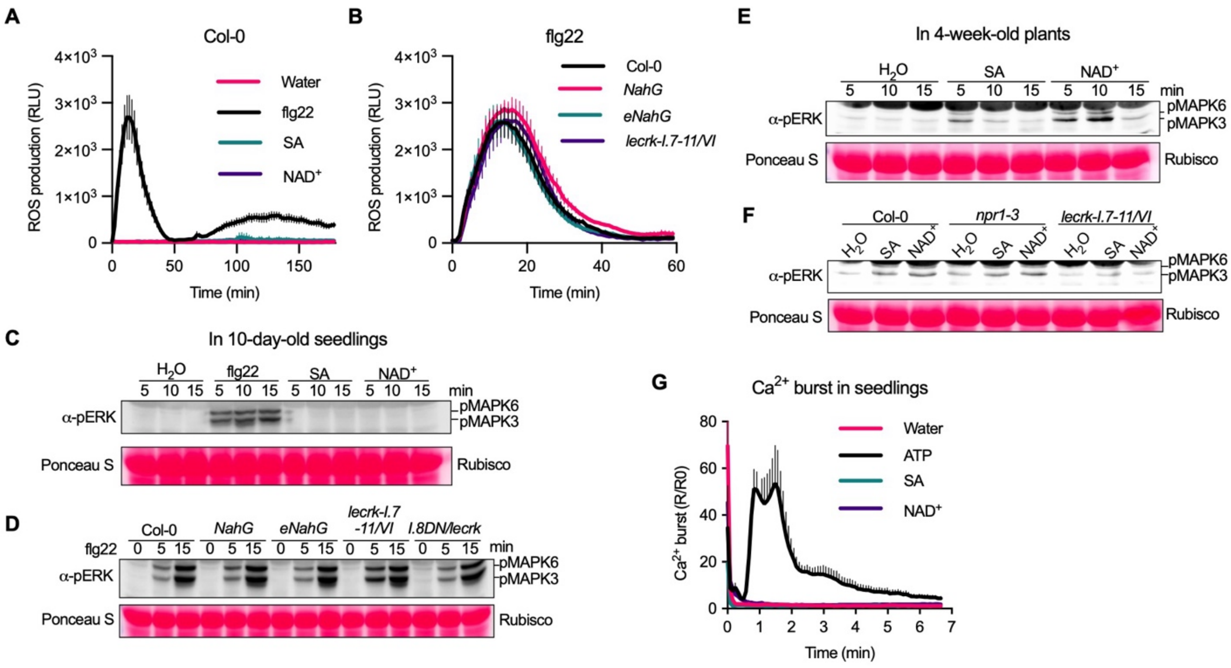


**Fig. S10.**

**SA- and NAD$^+$-induced early signaling events** (A) SA and NAD$^+$ did not trigger a detectable ROS burst. Leaf discs from 4-week-old Col-0 plants were treated with 0.5 mM SA or 0.5 mM NAD$^+$. Water and 100 nM flg22 served as negative and positive controls, respectively. (B) 100 nM flg22-induced ROS burst in *eNahG* and *lecrk* mutants. In (A and B), ROS production was recorded as RLU immediately after elicitor application for the indicated times, as described in Methods. Data represent means ± SEM (n = 8). (C) SA and NAD$^+$ failed to trigger MAPK activation in seedlings. Ten-day-old Col-0 seedlings were treated with 0.5 mM SA or 0.5 mM NAD$^+$ for indicated times. Water and 100 nM flg22 served as the negative and positive controls, respectively. (D) flg22-induced MAPK activation in *eNahG* and *lecrk* seedlings. Ten-day-old Col-0 seedlings were treated with 100 nM flg22 for indicated times. (E) SA- and NAD$^+$-triggered MAPK activation in adult plants. Leaf discs from 4-week-old Col-0 plants were treated with 0.5 mM SA or 0.5 mM NAD$^+$ for indicated times. Water served as the negative control. (F) SA- and NAD$^+$-induced MAPK activation in *npr1-3* and *lecrk-I.7-11/VI* adult plants. Leaf discs of 4-week-old plants were treated with water, 0.5 mM SA, or 0.5 mM NAD$^+$ for 5 min. In (C to F), treated seedling or leaf discs were subjected to MAPK activation assays as described in Methods. Ponceau S staining of Rubisco served as loading controls. (G) SA and NAD$^+$ failed to trigger a detectable cytosolic $Ca^{2+}$ burst in seedlings. Ten-day-old cytAEQ seedlings were treated with 0.5 mM SA or 0.5 mM NAD$^+$. Water and 0.2 mM ATP served as negative and positive controls, respectively. Cytosolic $Ca^{2+}$ burst was recorded as luminescence (R) immediately after treatment for indicated times, normalized to baseline luminescence before treatment ($R/R_0$). Data represent means ± SEM (n = 8).

### Table S1.
### Plant materials created in this study

| Material name | Methos used | Genes |
|---|---|---|
| *lecrk-I.1-6* | CRISPR/Cas9 | Clade I LecRK members 1-6 (6 genes) |
| *lecrk-I.7-11* | CRISPR/Cas9 | Clade I LecRK members 7-11 (5 genes) |
| *lecrk-II/III* | CRISPR/Cas9 | Clade II and III LecRKs (4 genes) |
| *lecrk-IV* | CRISPR/Cas9 | Clade IV LecRKs (4 genes) |
| *lecrk-V.1/2/9* | CRISPR/Cas9 | Clade V LecRK members 1, 2, and 9 (3 genes) |
| *lecrk-V.3/4* | CRISPR/Cas9 | Clade V LecRK members 3 and 4 (2 genes) |
| *lecrk-V.5/6/7/8* | CRISPR/Cas9 | Clade V LecRK members 5, 6, 7, and 8 (4 genes) |
| *lecrk-VII/VIII* | CRISPR/Cas9 | Clade VII and VIII LecRKs (4 genes) |
| *lecrk-IX* | CRISPR/Cas9 | Clade IX LecRKs (2 genes) |
| *lecrk-S.1/4/6/7* | CRISPR/Cas9 | Singleton 1, 4, 6, and 7 LecRKs (4 genes) |
| *lecrk-S.5* | CRISPR/Cas9 | Singleton 5 LecRK (1 gene) |
| *lecrk-I.8-cr* | CRISPR/Cas9 | LecRK-I.8 (1 gene) |
| *lecrk-I.1-6/VI* | Cross | Clade I LecRK members 1-6 and clade VI LecRKs (10 genes) |
| *lecrk-I.7-11/VI* | Cross | Clade I LecRK members 7-11 and clade VI LecRKs (9 genes) |
| *lecrk-I/VI* | Cross | Clade I and clade VI LecRKs (15 genes) |
| *35S:eNahG-GFP* | Transgenic | 35S:eNahG-GFP in Col-0 |
| *pUBQ10:lecrk-I.8-2DN*/Col-0 | Transgenic | pUBQ10:lecrk-I.8-2DN in Col-0 |
| *pUBQ10:lecrk-I.8-2DN/lecrk-VI* | Transgenic | pUBQ10:lecrk-I.8-2DN in lecrk-VI |
| *pUBQ10:lecrk-I.8-2DN/lecrk-I.8-2/VI (I.8DN/lecrk)* | Transgenic | pUBQ10:lecrk-I.8-2DN in lecrk-I.8-2/VI |
| *pUBQ10:LecRK-I.8-HA/lecrk-I.8-2* | Transgenic | pUBQ10:LecRK-I.8-HA in lecrk-I.8-2 |
| *pUBQ10:LecRK-I.8K357E-HA/lecrk-I.8-2* | Transgenic | pUBQ10:LecRK-I.8K357E-HA in lecrk-I.8-2 |
| *pUBQ10:LecRK-VI.2-HA/lecrk-VI.2-1* | Transgenic | pUBQ10:LecRK-VI.2-HA in lecrk-VI.2-1 |
| *pUBQ10:lecrk-VI.2$^{3A}$-HA/lecrk-VI.2-1* | Transgenic | pUBQ10:lecrk-VI.2$^{3A}$-HA in lecrk-VI.2-1 |
| *pUBQ10:lecrk-VI.2$^{\Delta loop}$-HA/lecrk-VI.2-1* | Transgenic | pUBQ10:lecrk-VI.2$^{\Delta loop}$-HA in lecrk-VI.2-1 |

**Table S2.**
**Primers used in this study**

| Name | Sequence (5’-3’) | Usage |
|---|---|---|
| *18KOBsaF* | ATATATGGTCTCGATTGCTCGAGATCCCAAACTGAGGTTTAAGAGCTATGCTGGAAACA | CRISPR vector |
| *18KOBsaR* | ATTATTGGTCTCGAAACTGGCGAAAACCGTTGAAGCTTGCACCAGCCGGGAATCG | CRISPR vector |
| *I.1-6KO-F* | TATGGTCTCGATTGTTGTTAGCAGAGGAAGGAGGTTTAAGAGCTATGCTGGAA | CRISPR vector |
| *I.1-6KO-R* | ATTGGTCTCGAAACATATCATCAACAACACACCCTGCACCAGCCGGGAATCG | CRISPR vector |
| *LRK1.7-1.11KOF* | TATGGTCTCGATTG TTTCATTACCAATGATTCG GTTTAAGAGCTATGCTG | CRISPR vector |
| *LRK1.7-1.11KOR* | ATTGGTCTCGAAAC TGGAGCAAGTGATACAATAC TGCACCAGCCGGGA | CRISPR vector |
| *II/III-KO-F1* | TATGGTCTCGATTGCACCACTGAAGTCCATCGAGTTTAAGAGCTATGCTGGAA | CRISPR vector |
| *II/III-KO-R1* | ATTGGTCTCGGACTTTAGCGGGGCTGCACCAGCCGGGAATCG | CRISPR vector |
| *II/III-KO-F2* | TATGGTCTCGAGTCCAGCGAGTTTAAGAGCTATGCTGGAA | CRISPR vector |
| *II/III-KO-R2* | ATTGGTCTCGAGCGGCAGACTTATTGCACCAGCCGGGAATCG | CRISPR vector |
| *II/III-KO-F3* | TATGGTCTCGCGCTTGCTAGGTTTAAGAGCTATGCTGGAA | CRISPR vector |
| *II/III-KO-R3* | ATTGGTCTCGAAACTTGCTAGGTGACGGTCAAGGTGCACCAGCCGGGAATCG | CRISPR vector |
| *IV.1-4KO-F1* | TATGGTCTCGATTGCTTATTCTATAGTCTCTGGGTTTAAGAGCTATGCTGGAA | CRISPR vector |
| *IV.1-4KO-R1* | ATTGGTCTCGACCGACAGACATATTGCACCAGCCGGGAATCG | CRISPR vector |
| *IV.1-4KO-F2* | TATGGTCTCGCGGTGGAGGAGTTTAAGAGCTATGCTGGAA | CRISPR vector |
| *IV.1-4KO-R2* | ATTGGTCTCGATCGCGATCTGAGATGCACCAGCCGGGAATCG | CRISPR vector |
| *IV.1-4KO-F3* | TATGGTCTCGCGATTCTCAGGTTTAAGAGCTATGCTGGAA | CRISPR vector |
| *IV.1-4KO-R3* | ATTGGTCTCGAAACATGTATTGACTAGGAAGTGCTGCACCAGCCGGGAATCG | CRISPR vector |
| *V.1/2/9-F1* | TATGGTCTCGATTGGAATATGTTGAGGAGCAAGGTTTAAGAGCTATGCTGGAA | CRISPR vector |
| *V.1/2/9-R1* | ATTGGTCTCGTTTAGATCCGCAACTGCACCAGCCGGGAATCG | CRISPR vector |
| *V.1/2/9-F2* | TATGGTCTCGTAAACCCGGAGTTTAAGAGCTATGCTGGAA | CRISPR vector |
| *V.1/2/9-R2* | ATTGGTCTCGAAACTAGATATTGTGAGGATGAACTGCACCAGCCGGGAATCG | CRISPR vector |
| *V.3/4-F1* | TATGGTCTCGATTGAAACCATTCATTAACCCGGGTTTAAGAGCTATGCTGGAA | CRISPR vector |
| *V.3/4-R1* | ATTGGTCTCGAAACTCACGAAGACAATAGAGAACTGCACCAGCCGGGAATCG | CRISPR vector |
| *V.5/6/7/8-F1* | TATGGTCTCGATTGTAAGGAACAGAACAAGAATGTTTAAGAGCTATGCTGGAA | CRISPR vector |
| *V.5/6/7/8-R1* | ATTGGTCTCGAAACTCCAGGGTCCAGTTTCCATTTGCACCAGCCGGGAATCG | CRISPR vector |
| *VII/VIII-KO-F1* | TATGGTCTCGATTGCACCCGGAAACTAAATCTCGTTTAAGAGCTATGCTGGAA | CRISPR vector |
| *VII/VIII-KO-R1* | ATTGGTCTCGTCAACGGTTCTAGCTGCACCAGCCGGGAATCG | CRISPR vector |
| *VII/VIII-KO-F2* | TATGGTCTCGTTGATTCCAGGTTTAAGAGCTATGCTGGAA | CRISPR vector |
| *VII/VIII-KO-R2* | ATTGGTCTCGTACCTTTCGCCACATGCACCAGCCGGGAATCG | CRISPR vector |
| *VII/VIII-KO-F3* | TATGGTCTCGGGTAACGGCGGTTTAAGAGCTATGCTGGAA | CRISPR vector |
| *VII/VIII-KO-R3* | ATTGGTCTCGAAACTTTGGACCGGTTAGACCAAGTGCACCAGCCGGGAATCG | CRISPR vector |
| *L9.1-2F1* | ATATATGGTCTCGATTGCAGGTAGTTGAGCTCCCATGTTTAAGAGCTAT | CRISPR vector |
| *L9.1-2R1* | ATTATTGGTCTCGCTTGCATCTCCTTGCACCAGCCGGGAATCG | CRISPR vector |
| *L9.1-2F2* | ATATATGGTCTCGCAAGAGCAAATGGTTTAAGAGCTATGCTGGAAACA | CRISPR vector |
| *L9.1-2R2* | ATTATTGGTCTCGCAACTCCCGATGTGCACCAGCCGGGAATCG | CRISPR vector |
| *L9.1-2F3* | ATATATGGTCTCGGTTGCATCCCCGGTTTAAGAGCTATGCTGGAAACA | CRISPR vector |
| *L9.1-2R3* | ATTATTGGTCTCGAAACCTCGATGTCAAATCCCATGTTGCACCAGCCG | CRISPR vector |
| *S.1/4/6/7-F1* | TATGGTCTCGATTGATACTTTAGGAAGTCCCGGGTTTAAGAGCTATGCTGGAA | CRISPR vector |
| *S.1/4/6/7-R1* | ATTGGTCTCGCGTCCTCTCCAATTTGCACCAGCCGGGAATCG | CRISPR vector |
| *S.1/4/6/7-F2* | TATGGTCTCGGACGAAGCAGGTTTAAGAGCTATGCTGGAA | CRISPR vector |
| *S.1/4/6/7-R2* | ATTGGTCTCGTCTGGAGTTGGAGTTGCACCAGCCGGGAATCG | CRISPR vector |
| *S.1/4/6/7-F3* | TATGGTCTCGCAGATCTCAGGTTTAAGAGCTATGCTGGAA | CRISPR vector |
| *S.1/4/6/7-R3* | ATTGGTCTCGAAACCTTCTTTCTCTTGCCGCTTCTGCACCAGCCGGGAATCG | CRISPR vector |
| *S.2/3/5-F1* | TATGGTCTCGATTGTAACAACGCCTTGTAAACGGTTTAAGAGCTATGCTGGAA | CRISPR vector |
| *S.2/3/5-R1* | ATTGGTCTCGTGCTTTGTAGAGAGTGCACCAGCCGGGAATCG | CRISPR vector |
| *S.2/3/5-F2* | TATGGTCTCGAGCAACCAAAGTTTAAGAGCTATGCTGGAA | CRISPR vector |
| *S.2/3/5-R2* | ATTGGTCTCGAAACCCGGGATCGGTTTTGTTAGATGCACCAGCCGGGAATCG | CRISPR vector |
| *I.1-6KO-gtF* | TCATAAGAGCCCTGATTTGTTATT | CRISPR genotyping |
| *I.1-6KO-gtR* | AGTGTTCATATTTGGTGAGAACAT | CRISPR genotyping |
| *LRK1.7-1.11KOgtF* | GTCGTATTCAATGGGTCTACAGC | CRISPR genotyping |
| *LRK1.7-1.11KOgtR* | TCACCTCCCATCTCCGTATGT | CRISPR genotyping |
| *II.1genoF* | GTGGTATCTCCAACCAAAGATCT | CRISPR genotyping |
| *II.1genoR* | GCTTGAAACTCCATCCAAGGATA | CRISPR genotyping |
| *II.2genoF* | TGATAATATGTGTCCAGGTGTTG | CRISPR genotyping |
| *II.2genoR* | GTACCGCTAAAGTAACTAGCATT | CRISPR genotyping |
| *III.1genoF* | AGTTTCTTAACCATGGCTTCCTC | CRISPR genotyping |
| *III.1genoR* | CTTACTGGTTTCCCACTCGCC | CRISPR genotyping |
| *III.2genoF* | CCAAATTTCTTAACCATGGCTTT | CRISPR genotyping |
| *III.2genoR* | CTTATTGGTTTCCCACTCGCC | CRISPR genotyping |
| *L4.1genoF* | TTTGCTTACAACAATGGCTTTAAT | CRISPR genotyping |
| *L4.1genoR* | AATCGACCCAGACCTGCATCGGC | CRISPR genotyping |
| *L4.2genoF* | CACCTACAATGGTTTCCATCCTC | CRISPR genotyping |
| *L4.2genoR* | GAGCCACGGTGACATCAATTCGA | CRISPR genotyping |
| *L4.34genoF* | AGGTAACCCTCGATTGGAAACAGAG | CRISPR genotyping |
| *L4.34genoR* | TCACCTCCCACTGGAGAGTAGAG | CRISPR genotyping |
| *V.12gF1* | CTCCTGTCGCGCCGAATATAG | CRISPR genotyping |
| *V.12gR1* | CGAACCGCAACATGGTTATATCG | CRISPR genotyping |
| *V.12gF2* | GAGTCTGTAGACTATAAACGTGAGG | CRISPR genotyping |
| *V.12gR2* | ATTTGGAGCTATCATTCTTGGTTTC | CRISPR genotyping |
| *V.9F1* | TAGTGCTTCTTCTGGTGCTGCA | CRISPR genotyping |
| *V.9R1* | GGTTAGAGAAAGCAATGGAATCTT | CRISPR genotyping |
| *V.3gF* | GGAACATTCGGGTACATGGCG | CRISPR genotyping |
| *V.3gR* | CTAGAGAAACAACCTGGGTCTGC | CRISPR genotyping |
| *V.34gF* | CATGCCACTGCCAACTCTATGGC | CRISPR genotyping |
| *V.34gR* | CCGAACTCAGCATCACGTCTCA | CRISPR genotyping |

| | | |
|---|---|---|
| *V.5678gF* | GTTTGATAAGTTATGTTCCACGAAA | CRISPR genotyping |
| *V.5678gR* | GCTTTACGACTTGGTTTAGGCC | CRISPR genotyping |
| *VII.1gF* | CAACGGCTTCAATGACTCATCAT | CRISPR genotyping |
| *VII.1gR* | CCATCGTTAAGCTTCAATGGCTT | CRISPR genotyping |
| *VII.2gF* | CTCGCATCCCTACTCTTGTTCCG | CRISPR genotyping |
| *VII.2gR* | GTGGTTGTCGTTGATGTCGTTGA | CRISPR genotyping |
| *VIII.1gF* | GCAATGGAATCGTTGGCCTCACA | CRISPR genotyping |
| *VIII.1gR* | GAGACTGATACGTTGAAGACCCG | CRISPR genotyping |
| *VIII.2gF* | CTCAACTCTCATCCACATTCTTG | CRISPR genotyping |
| *VIII.2gR* | GATCCAATCCCACGTGGTTCCC | CRISPR genotyping |
| *L9.1genoF* | GCCTGCAAATGGCCAACTCAAT | CRISPR genotyping |
| *L9.1genoR* | TATTTCCCTCGGTGACCCCTCC | CRISPR genotyping |
| *L9.2genoF* | CCTCCTCTATGTCTAATTCAATC | CRISPR genotyping |
| *L9.2genoR* | CTTTGCATGACAAATATCTTGGC | CRISPR genotyping |
| *S1gF* | TCTCTACAATTCCTTCAGCTCCG | CRISPR genotyping |
| *S1gR* | CGCATATTAAAGCGAACGAAGCT | CRISPR genotyping |
| *S4gF* | GCCTCACCAAATTTAACGTTAAAC | CRISPR genotyping |
| *S4gR* | GAATCGTAATCAATCCAAGCTTG | CRISPR genotyping |
| *S6gF* | CGTCATCTTCCACCTAATTCTCTTC | CRISPR genotyping |
| *S6gR* | CAACGGAAGAAACAGAGAAGATT | CRISPR genotyping |
| *S7gF* | CATTCCTCGGAGATTCACATCTC | CRISPR genotyping |
| *S7gR* | GAACACGTTCAATAAACGTAAATCG | CRISPR genotyping |
| *S.5F1* | CTCTTGCGTGGAAGCTCCTGTT | CRISPR genotyping |
| *S.5R1* | GCTTGTTGAAGCTGTAAATCCAAC | CRISPR genotyping |
| *SK072930F* | TGTGTCATCCATATCTTGCAATG | T-DNA genotyping |
| *SK072930R* | AGATGTGTTTGCAGAGTGACAAG | T-DNA genotyping |
| *SK026551F* | GACCCAAACACACACTTCTCTTC | T-DNA genotyping |
| *SK026551R* | TGCATATCCAGATTTACATCTTTTTG | T-DNA genotyping |
| *G20-I.8-2F* | CCGTCGACCTCGAGGAAAATGGCTCCAGGATTGGATCTGATC | Overexpression vector |
| *G20-I.8-2R* | TCCTACCCGGGAGCGTCAAATTTCAGCTACGAATTGCTTCAT | Overexpression vector |
| *seq-NahG* | CCAACCCGCGAAGGCATCGA | Overexpression vector |
| *SacI-SPF* | CGAGCTCATGAATTTTACTGGCTATTCTCGATTTTTAATCGTCTTTGTAGCTCTT | Overexpression vector |
| *NahGF1* | TGTAGCTCTTGTAGGTGCTCTTGTTCTTCCCTCGAAAGCTCAAGATAAAAAC | Overexpression vector |
| *XbaI-NahGR* | GCTCTAGACCCTTGACGTAGCGCAC | Overexpression vector |
| *SalI-AHA2F* | CTCTAGAGGTGTCGACATGTCGAGTCTCGAAGATATCAAG | Transient expression vector |
| *SalI-AHA2R* | CTCTAGAGGTGTCGACCACAGTGTAGTGACTGGGAGTT | Transient expression vector |
| *HREe1.8F* | TAAACGTCTCTAAAAATGGCTCCAGGATTGGATCTGA | Transient expression vector |
| *HREe1.8R* | AATGAAACCAGAGCGTCAGTGATGGTGATGGTGATGAGAAGTTTTCATTTTAGGATGAG | Transient expression vector |
| *HREe6.2F* | TATGGTCTCGAAAATGGGCACACAAAGATCCAT | Transient expression vector |
| *HREe6.2HisR* | TATGGTCTCTAGCGTCAGTGATGGTGATGGTGATGAGATCCTTCCTTCTTCGCCTTATTCGG | Transient expression vector |
| *HREeV.5F* | TAAACGTCTCTAAAAATGTCTCGTGAACTTATTATTCTCTG | Transient expression vector |
| *HREeV.5R* | AATGAAACCAGAGCGTCAGTGATGGTGATGGTGATGGGATCCAGATGTTTTCTTTGGATATGGAGG | Transient expression vector |
| *HREeBAK1F* | TAAACGTCTCTAAAAATGGAACGAAGATTAATGATCCCT | Transient expression vector |
| *HREeBAK1R* | AATGAAACCAGAGCGTCAGTGATGGTGATGGTGATGACTCCCTGCAGGTGATGGCG | Transient expression vector |
| *NahG-HisF* | AGGAGATATACCATGGATGAAAAACAATAAACTTGGCTTGC | Protein expression vector |
| *NahG-HisR* | CCGCAAGCTTGTCGACCCCTTGACGTAGCGCACCCC | Protein expression vector |
| *UBQ10_1.8_Kpn1F* | CCGTCGACCTCGAGGATGGCTCCAGGATTGGATCTGA | Complementation vector |
| *L18HA-R* | ACATCATAAGGGTAGGTACCTCGTCCAATTCCGTATTGAAT | Complementation vector |
| *L18K366E-R* | AATCTTTCCACAGCTATGTCTTCCTGAG | Complementation vector |
| *L18K366E-F* | AGCTGTGGAAAGATTTTCCCATCATGGA | Complementation vector |
| *VI.2de12F* | TCGACACGGGAAACGACATCGGTCTGAA | Complementation vector |
| *VI.2de12R* | CGTTTCCCGTGTCGAATTCCACTGCGA | Complementation vector |
| *VI.2F89AF* | AAAGGCGCTGGATTCACATTCACACTATCTC | Complementation vector |
| *VI.2F89SR* | GAATCCAGCGCCTTTATTGCTTGAGCTTGAA | Complementation vector |
| *VI.2R143AR* | ATGTCGTTTCCTATTGCGTCGGTGTTGTCGTCTCTTG | Complementation vector |
| *VI.2R155R* | GCACTGTTGTAATTCAGACCGATGTCGTTTCCTATTCTGTC | Complementation vector |
| *VI.2R155A* | GAATTACAACAGTGCTACTTCGGATCTCCAAGAACCA | Complementation vector |
| *KpnI-AEQ-F* | CCGTCGACCTCGAGGGTACCATGACCAGCGAACAATACTCAGT | Protoplast expression |
| *KpnI-AEQ-R* | TCCTACCCGGGAGCGTTAGGGGACAGCTCCACCGTA | Protoplast expression |
| *PR1-F* | CTCATACACTCTGGTGGG | qPCR |
| *PR1-R* | ATTGCACGTGTTCGCAGC | qPCR |
| *UBQ5-F* | TCTCCGTGGTGGTGCTAAG | qPCR |
| *UBQ5-R* | GAACCTTTCCAGATCCATCG | qPCR |
| *qPR2F* | ATCAAGGAGCTTAGCCTCAC | qPCR |
| *qPR2R* | TGTAAAGAGCCACAACGTCC | qPCR |
| *qPR5F* | CTCTTCCTCGTGTTCATCAC | qPCR |
| *qPR5R* | GAAGCACCTGGAGTCAATTC | qPCR |
| *qUGT76B1-F* | TGGAAGATCGGATTGCATT | qPCR |
| *qUGT76B1-R* | CCTTCATGGGCATAATCCTC | qPCR |
| *qFMO1-F* | CTTGGCTTGAGTTTCCAAGC | qPCR |
| *qFMO1-R* | CCACATTGAACGTAGCTCTG | qPCR |
| *qALD1-F* | GGATTGGCATGCCTTTCTTC | qPCR |
| *qALD1-R* | TGAACCCACAAGTATGGAGC | qPCR |
| *Q-WRKY18F* | TTAGATGCTCGTTTGCACCG | qPCR |
| *Q-WRKY18R* | CCAAAGTCACTGTGCTTGAC | qPCR |
| *Q-WRKY38F* | GAATTGGAGGGACGATTCAC | qPCR |
| *Q-WRKY38R* | CATGCTTTCTTGCTTCGCAG | qPCR |

**Data S1. (separate file)**
RNA-seq data of SA-induced transcriptional reprogramming in Col-0, *I.7-11/VI*, and *bak1-5 bkk1*

**Data S2. (separate file)**
RNA-seq data of SA-induced transcriptional reprogramming in Col-0 and *I.8DN/lecrk*

**Data S3. (separate file)**
RNA-seq data of *Psm*-induced transcriptional reprogramming in Col-0, *NahG*, and *eNahG*

**Data S4. (separate file)**
Phosphoproteomic data of SA-induced phosphorylation in Col-0, *I.7-11/VI*, and *bak1-5 bkk1*
